# Evolution of an interaction between disordered proteins resulted in increased heterogeneity of the binding transition state

**DOI:** 10.1101/2020.03.27.012328

**Authors:** Elin Karlsson, Cristina Paissoni, Amanda M. Erkelens, Zeinab Amiri Tehranizadeh, Frieda A. Sorgenfrei, Eva Andersson, Weihua Ye, Carlo Camilloni, Per Jemth

## Abstract

Intrinsically disordered protein (IDP) domains often have multiple binding partners. Little is known regarding molecular changes in the binding mechanism when a new interaction evolves from low to high affinity. Here we compared the degree of native contacts in the transition state of the interaction of two IDP domains, low-affinity ancestral and high-affinity human NCBD and CID. We found that the coupled binding and folding mechanism of the domains is overall similar, but with a higher degree of native hydrophobic contact formation in the transition state of the ancestral complex while more heterogenous transient interactions, including electrostatic, and an increased disorder characterize the human complex. From an evolutionary perspective, adaptation to new binding partners for IDPs may benefit from this ability to exploit multiple alternative transient interactions while retaining the overall pathway.

## Introduction

Intrinsically disordered proteins (IDPs) are abundant in the human proteome^1^ and are frequently involved in mediating protein-protein interactions in the cell. The functional advantages of disordered proteins include exposure of linear motifs for association with other proteins, accessibility for post-translational modifications, formation of large binding interfaces and the ability to interact specifically with multiple partners. These properties make IDPs suitable for regulatory functions in the cell, wherefore IDPs often act as hubs in interaction networks governing signal transduction pathways and transcriptional regulation^2^.

Despite recent appreciation of the biological importance of IDPs and progress in understanding their mechanism of interaction with other proteins, the changes that take place at a molecular level when these proteins evolve to bind new binding partners remain elusive. The reason for this is the inherent difficulty in assessing effects of mutations that a protein has acquired during millions or billions of years. However, the rapidly increasing number of available protein sequences from extant species has enabled the development of ancestral sequence reconstruction as a tool for inferring the evolutionary history of proteins^3^. Ancestral sequence reconstruction relies on an alignment of sequences from extant species and a maximum likelihood method that infers probabilities for amino acids at each position in the reconstructed ancestral protein from a common ancestor. However, due to less constraints for maintaining a folded structure, IDPs in general experience faster amino acid substitution rates during evolution and an increased occurrence of amino acid deletion and insertion events as compared to folded proteins, which often obstructs reliable sequence alignments of IDPs^4–6^.

Nevertheless, IDP regions that form binding interfaces in coupled binding and folding interactions are usually conserved because of sequence restraints for maintaining affinity and structure of the protein complex^7^, and can therefore be subjected to ancestral sequence reconstruction. The ability to mutate while maintaining certain sequence characteristics might allow IDPs to more efficiently explore sequence space, facilitating adaptation to new binding partners.

The nuclear co-activator binding domain (NCBD) from CREB-binding protein (CBP) is engaged in multiple protein-protein interactions in the cell^8^. NCBD is a molten globule-like protein^9^, which forms three helices in the unbound state that rearranges upon binding to its different partners^10–12^. These binding partners include among others the transcription factors and transcriptional co-regulators p53, IRF3 and NCOA1, 2 and 3 (also called SRC, TIF2 and ACTR, respectively). The interaction between NCBD and the CBP-interacting domain (CID) from NCOA3 (ACTR) has been intensively studied with kinetic methods to elucidate details about the binding mechanism^13–16^. These protein domains interact in a coupled folding and binding reaction in which CID forms two to three helices that wrap around NCBD^10^. The binding reaction involves several steps, as evidenced from the multiple kinetic phases observed in stopped flow spectroscopy and single molecule-FRET experiments^16–18^. It is however not clear what all kinetic phases corresponds to and equilibrium data agree well with the major kinetic binding phase^19^ in overall agreement with a two-state binding mechanism.

Previous phylogenetic analyses revealed that while NCBD was present already in the last common ancestor of all bilaterian animals (deuterostomes and protostomes) CID likely emerged later as an interaction domain within the ancestral ACTR/TIF2/SRC protein in the deuterostome lineage (including chordates, echinoderms and hemichordates; Figure 1)^20^. The evolution of the NCBD/CID interaction was previously examined using ancestral sequence reconstruction in combination with several biophysical methods to assess differences in affinity, structure and dynamics between the extant human and ancestral protein complex^20, 21^. After the emergence of CID in the ancestral ACTR/TIF2/SRC, NCBD increased its affinity for CID while maintaining the affinity for some of its other binding partners^20^. While the overall structure of the ancestral NCBD/CID complex, which we denote as “Cambrian-like”, is similar to the modern high affinity human one, there are several differences on the molecular level (Figure 1)^21^.

**Figure 1.**
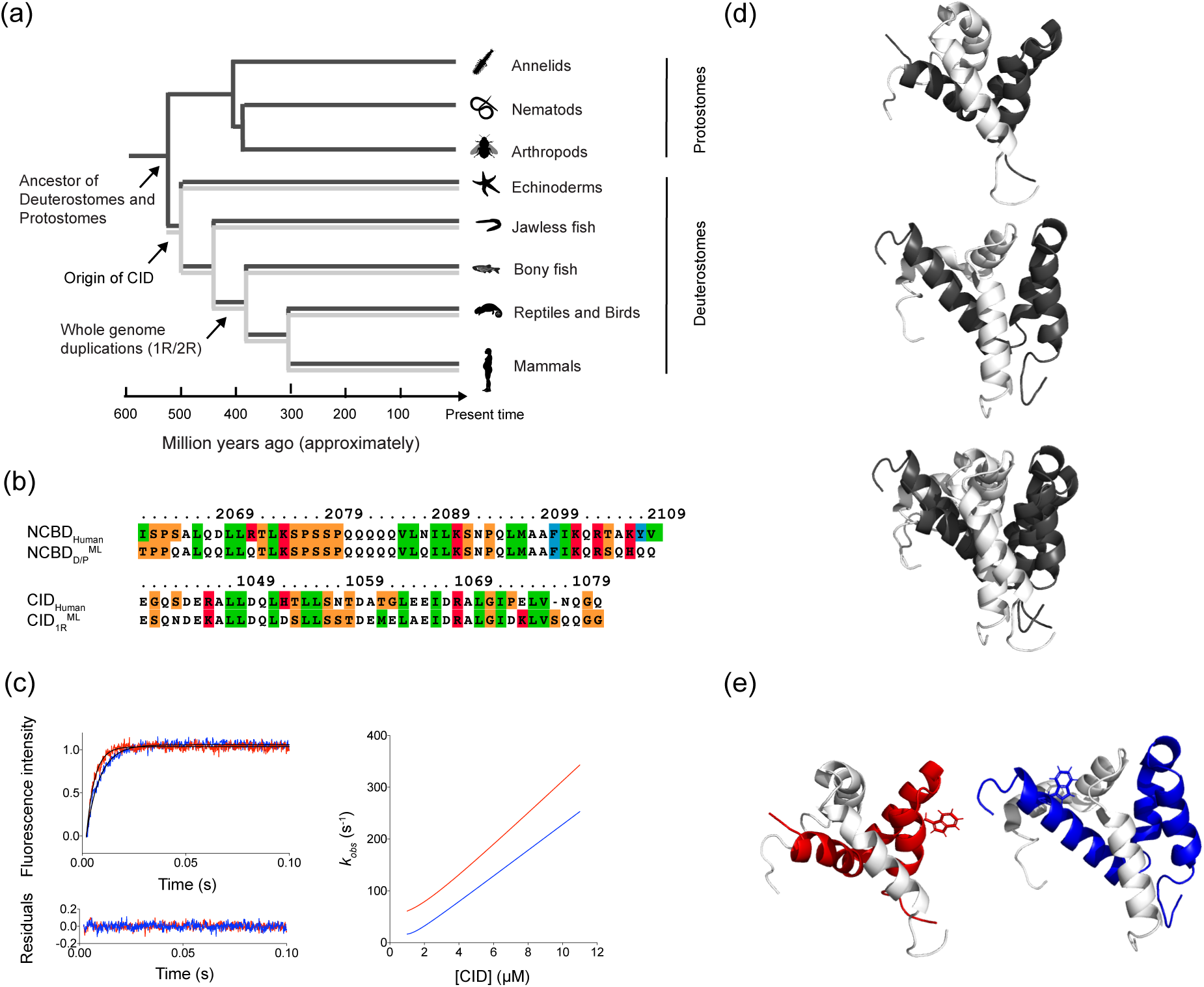
Extant and ancestral NCBD and CID variants. (a) A schematic phylogenetic tree showing the evolutionary relationship between extant species and reconstructed ancestral NCBD (dark grey) and CID (light grey) variants. The schematic animals were obtained from phylopic.org. (b) The sequences of the reconstructed ancestral and extant human NCBD (top) and CID (bottom) that have been used in the study. Human NCBD is from CREBBP and human CID is from NCOA3/ACTR. The color denotes residue type. (c) Examples of typical stopped-flow kinetic traces for the Cambrian-like complex (red) and human complex (blue) to the left. The concentrations used in this example were 1 µM NCBD and 6 µM CID for both experiments. The kinetic traces were fitted to a single exponential function (shown as a solid black line) and the residuals are displayed below the curve. The figure to the right shows the dependence of the observed rate constant (*kobs*) on CID concentration for the Cambrian-like complex (red) and the human complex (blue), using the rate constants obtained in global fitting (Table 2). (d) Solution structures of the Cambrian-like complex (top; PDB entry 6ES5)^31^, the human complex (middle; PDB entry 1KBH)^10^ and an alignment of the two complexes (bottom) with NCBD in dark grey and CID in light grey. (e) Structures of the Cambrian-like complex (left; NCBD in red and CID in light grey) and the extant human complex (right; NCBD in blue and CID in light grey) showing the position of the engineered Trp residues as stick model.

Here, we address the molecular details of the evolutionary optimization of the interaction between NCBD and CID. In order to investigate how evolution has modulated the binding energy landscape of this interaction, and in particular its transition state, we subjected the Cambrian-like NCBD/CID complex to site-directed mutagenesis and kinetic measurements using fluorescence-monitored stopped flow spectroscopy. The Cambrian-like complex consisted of the maximum likelihood estimate (ML) NCBD variant from the time of the divergence between deuterostomes and protostomes, NCBDD/P^ML^, and the CID variant from the time of the first whole genome duplication, CID1R. The experimental data were used to estimate the degree of native intermolecular tertiary contacts as well as helical content of CID in the transition state, expressed as phi (*ϕ*)-values. The *ϕ*-value, which commonly ranges from 0 to 1, reports on formation of native contacts in the transition state and was originally developed for folding studies^22^, however, it has also been used to gain detailed information about the transition state structures for several IDP binding reactions^23^. Furthermore, to obtain an atomic-level detail of the binding and folding process, the *ϕ*-values were employed for restrained molecular dynamics (MD) simulations. Our results suggest that while the overall binding mechanism and transition state are conserved, some modulation of helical content in the transition state and, most notably, an increased heterogeneity of transient intermolecular interactions is observed between the low-affinity Cambrian-like and high-affinity modern human complexes.

## Results

### Experimental strategy

We have characterized the transition state of binding for the low-affinity Cambrian-like NCBD/CID complex in terms of *ϕ*-values and compared it to that of the high-affinity modern human complex. A *ϕ*-value can be employed as a region-specific probe for native contact formation in the transition state of a binding reaction. A *ϕ*-value of 0 means that the entire effect on *K_D_* from the mutation stems from a change in *k_off_*, indicating that the interactions with the mutated residue mainly forms after the transition state barrier of binding. On the other hand, a *ϕ*-value of 1 is obtained when the effect of mutation on *k_on_* and *K_D_* is equally large, which suggests that the mutated residue is making fully formed native contacts at the top of the transition state barrier. To characterize the transition state for binding, we performed extensive site-directed mutagenesis in the Cambrian-like complex and subjected each mutant to stopped-flow kinetic experiments to obtain binding rate constants (i.e., *k_on_* and *k_off_*) (Figure 1 c). These rate constants were used to compute *ϕ*-values for each mutated position in the complex (Figure 2 a). To provide more structural details about the evolution of the NCBD/CID interaction, we also determined the TS ensemble of the Cambrian-like complex via *ϕ*-value-restrained MD simulations, following the same procedure previously used to obtain the TS ensemble of the human complex^24^.

**Figure 2.**
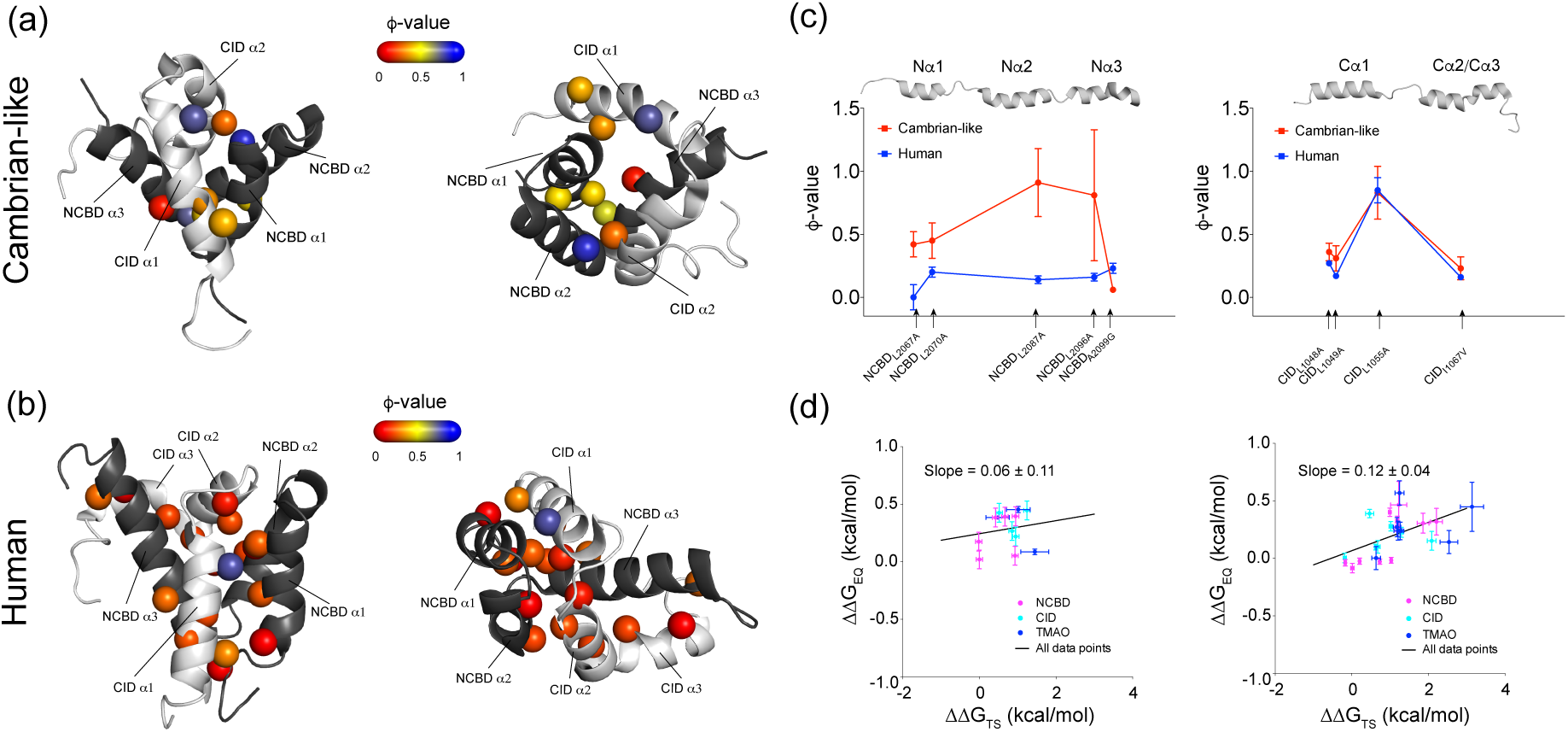
ϕ-values mapped onto the structures of the Cambrian-like and human complexes. (a) *ϕ*-values for conservative deletion mutations (mostly Leu*→*Ala mutations) in the binding interface of the Cambrian-like complex. NCBD in dark grey and CID in light grey. The two structures represent the same complex from different angles. Most *ϕ*-values fall within the intermediate to high *ϕ*-value category (0.3-0.9). (b) The previously published *ϕ*-values for conservative deletion mutations in the binding interface of the human NCBD/CID complex^13^. Most *ϕ*-values are in the low region (<0.3). (c) A site-to-site comparison between *ϕ*-values at corresponding positions in the Cambrian-like (red) and human (blue) complexes. (d) Brønsted plots for the Cambrian-like (left) and human (right) NCBD/CID interaction. Data for human NCBD/CID were obtained from previous studies^13, 24^. All structures were created using PyMOL.

In the present study we used NCBD_D/P_ as a pseudo wild type (Figure 1 e; denoted NCBD_D/P_) to obtain a sufficient signal change in the stopped flow fluorescence measurements. While Trp pseudo wild types have been shown to be reliable in protein folding studies^25^ it is less clear how much an engineered Trp would affect an IDP complex in a coupled binding and folding reaction. Therefore, we initially tested five different Trp mutants of NCBD_D/P_^ML^ to select the one with properties most similar to NCBD_D/P_^ML^ (Fig. S1). To check how robust our *ϕ*-values were to the position of the Trp probe we measured five *ϕ*-values with one of the other Trp variants, NCBD_D/P_. Despite a two-fold lower *k*_on_ as compared to NCBD_D/P_ (*i.e*., NCBD_D/P_) all five values were very similar, including the negative *ϕ*-value obtained for D1068A (Table 3), suggesting that our *ϕ*-values are robust and not dependent on the optical probe.

We constructed two sets of mutants in this study to characterize the transition state for binding: one that targeted native contacts in the binding interface and a second that targeted native helices in CID. The first set consisted of 13 NCBD_D/P_ variants and 10 CID_1R_ variants mainly corresponding to previously mutated residues in the human complex^13, 26^. The positions spanned the entire binding interface of NCBD and CID to ensure that all regions of the protein domains were probed (Table 1). The majority of the mutations targeted interactions between hydrophobic residues in the binding interface, however, a few mutations also probed interactions between charged residues. The second set of mutants consisted of Ala*→*Gly substitutions in helix 1 and 2 of CID_1R_, as well as helix 2 and 3 of CID_Human_ complementing a previous data set for helix-modulating mutations in helix 1 of CID_Human_^27^.

The secondary structure content of all mutants was assessed with far-UV CD. All CID variants exhibited far-UV CD spectra typical for highly disordered proteins (Figure S2 a-b). For NCBD, the far-UV CD measurements showed that NCBD_D/P_^L2070A^, NCBD_D/P_, NCBD_D/P_ and NCBD_D/P_ displayed substantially less *α*-helical structure than NCBD_D/P_, as judged from the lower magnitude of the CD signal at 222 nm (Figure S2 c). Addition of 0.7 M trimethylamine N-oxide (TMAO) to the experimental buffer for these variants resulted in an increase in helical content such that the far-UV CD spectra of NCBD_D/P_^L2070A^ and NCBD_D/P_^L2096A^ were qualitatively similar to NCBD_D/P_^pWT^ (Figure S2 d). For NCBD_D/P_^L2096A^ and NCBD_D/P_^A2099G^ stopped-flow kinetic experiments were carried out both in regular buffer and in buffer supplemented with 0.7 M TMAO. The resulting *ϕ*-values were very similar for these variants at both conditions (Figure S4). The complex of the NCBD_D/P_ variant was too unstable in buffer and data was only recorded in presence of 0.7 M TMAO. Several mutants failed to generate kinetic data due to elevated *k_obs_* values, which were too high to be reliably recorded with the stopped-flow technique. This was the case for NCBD_D/P_ ^L2074A^, NCBD_D/P_^L2086A^, NCBD_D/P_^L2090A^, NCBD_D/P_^I2101A^, CID_1R_^L1056A^, CID_1R_^L1064A^, CID_1R_^I1067A^ and CID_1R_^L1071A^.

### HSQC spectra suggest that the structure of the complex is robust to mutation

One caveat with *ϕ*-value analysis is the assumption that ground states are not affected by mutation. Hence, the strategy of using conservative deletion mutations is important^28^. For example, mutation to a larger residue is not considered conservative and one of the Trp variants that we tested, NCBDD/P, displayed a very low *k*on (3.5 μM s) and a 2-fold lower *k*off than the other NCBD variants (Fig. S1). We can only speculate about the structural basis for this result but since the residue is situated in the loop between N*α*1 and N*α*2 it might either lock the two helices in relation to each other or flip over, cover the binding groove and thus block access for CID. In either case we have a clear effect on the ground state. The problem of ground state changes may be particularly pertaining for IDPs, since they are more malleable in terms of structural changes to accommodate binding partners. It is beyond the scope of this study to perform extensive structure determination of all site-directed mutants but we recorded HSQC spectra for one complex containing a typical deletion mutation and an intermediate *ϕ*-value, NCBDD/P with CID1R (Fig. S3 a-b). The similar distribution of peaks, in particular for CID1R^ML^, suggests that the structure of the complex between CID1R^ML^ on the one hand and NCBDD/P^L2067A^ or NCBDD/P^pWT^ on the other are very similar.

### The transition state of the Cambrian-like complex is more native-like than the extant human complex

First, we assessed interactions formed by hydrophobic side chains in the binding interface of the ancestral Cambrian-like complex by computing *ϕ*-values for each mutant (Table 1). The resulting *ϕ*-values were mainly in the intermediate category ranging between 0.3-0.6 (Figure 2 a). This result was in contrast with previously published low *ϕ*-values for the human complex^29^, which ranged between 0-0.3 for similar conservative deletion mutations (Figure 2 b). A notable exception was the CIDHuman^L1055A^ variant, which displayed a *ϕ*-value of 0.83.

**Table 1.**
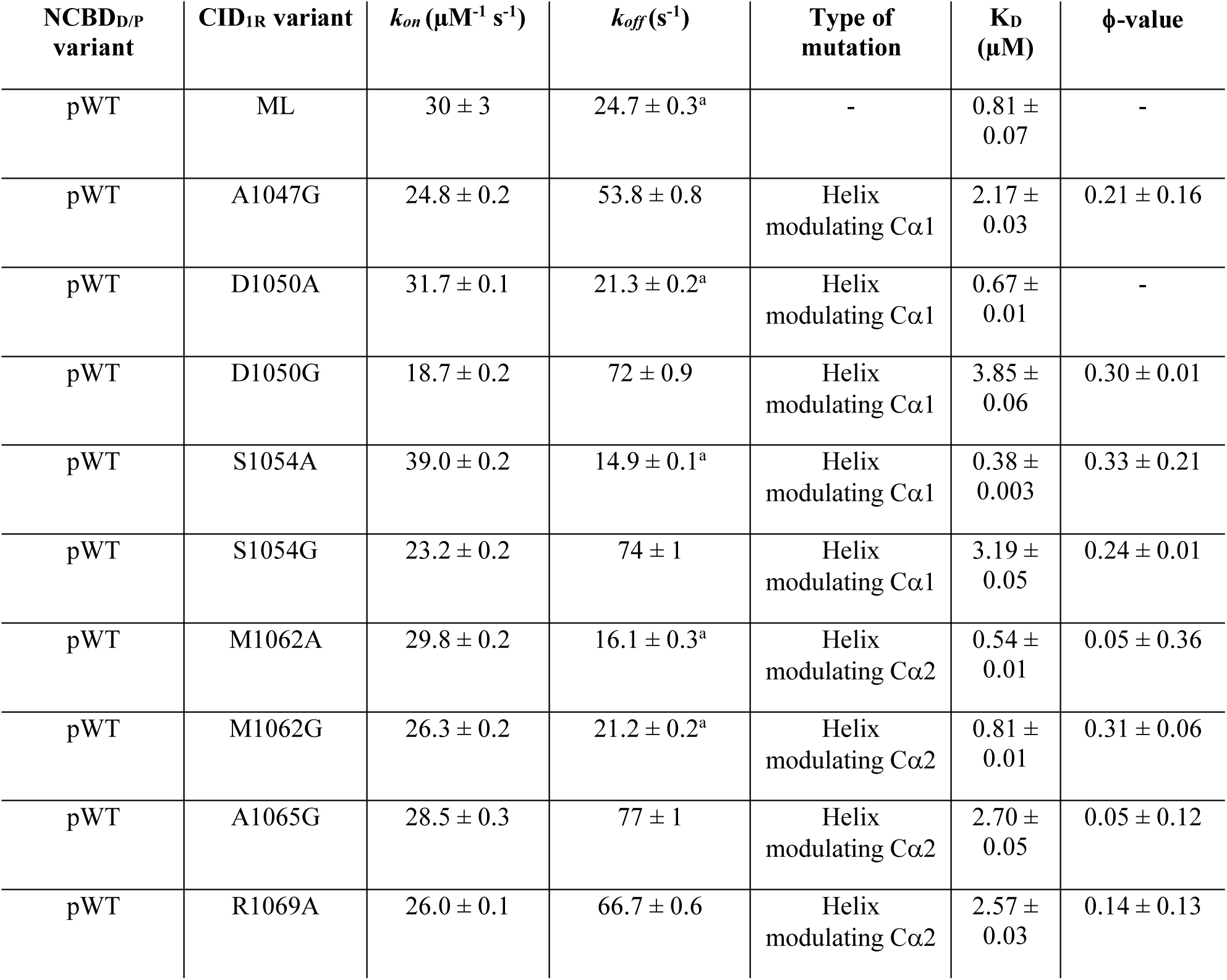
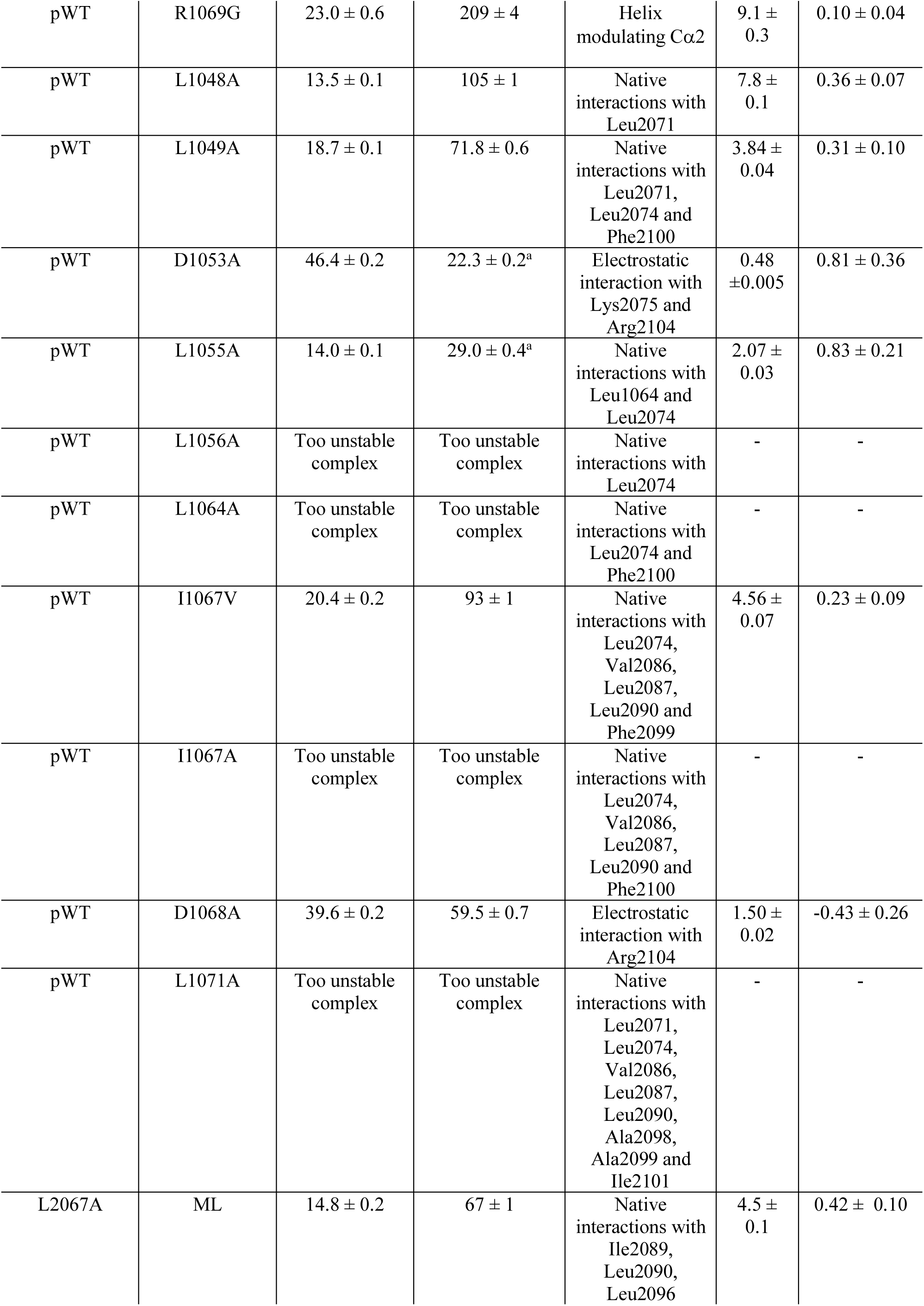
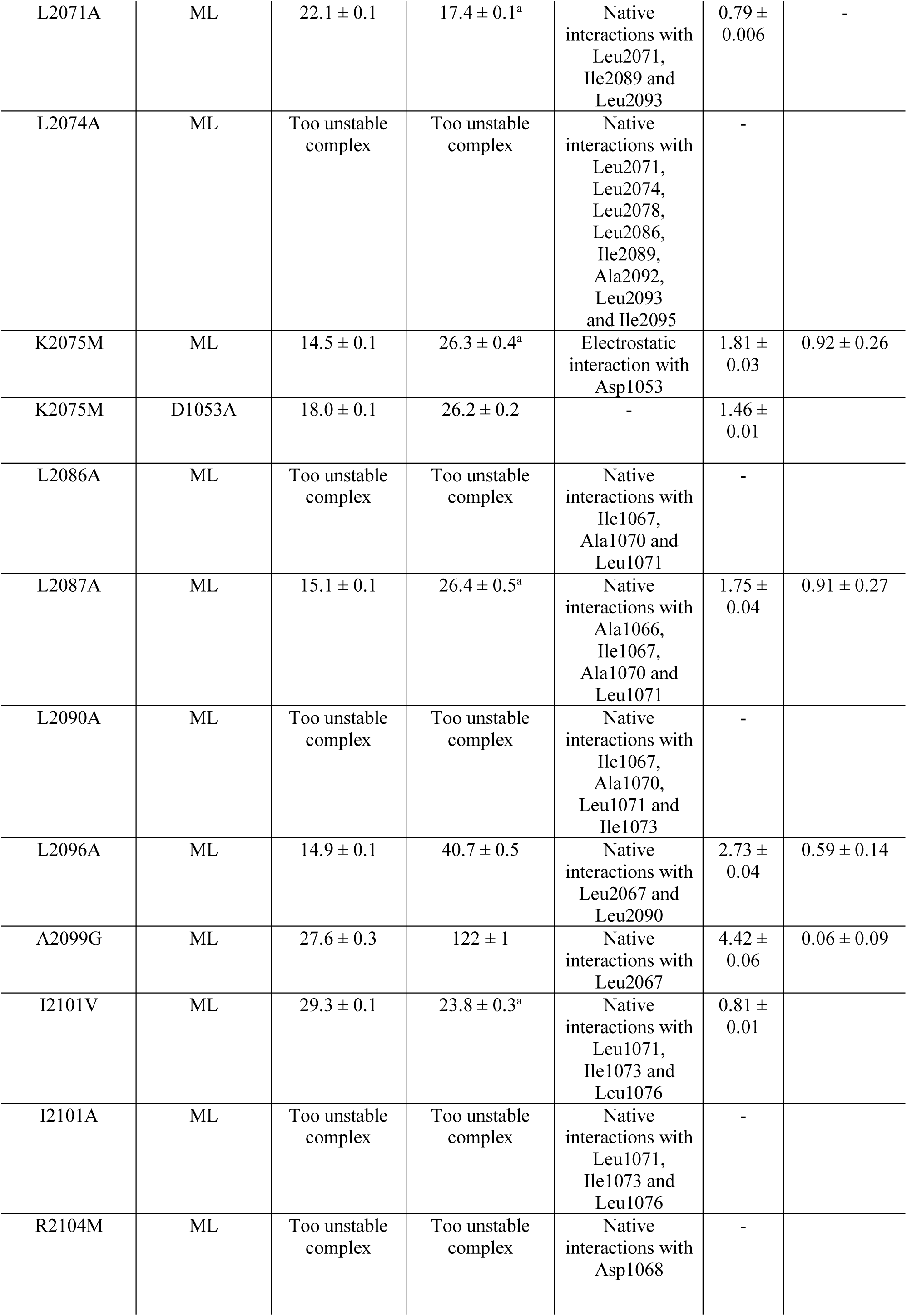
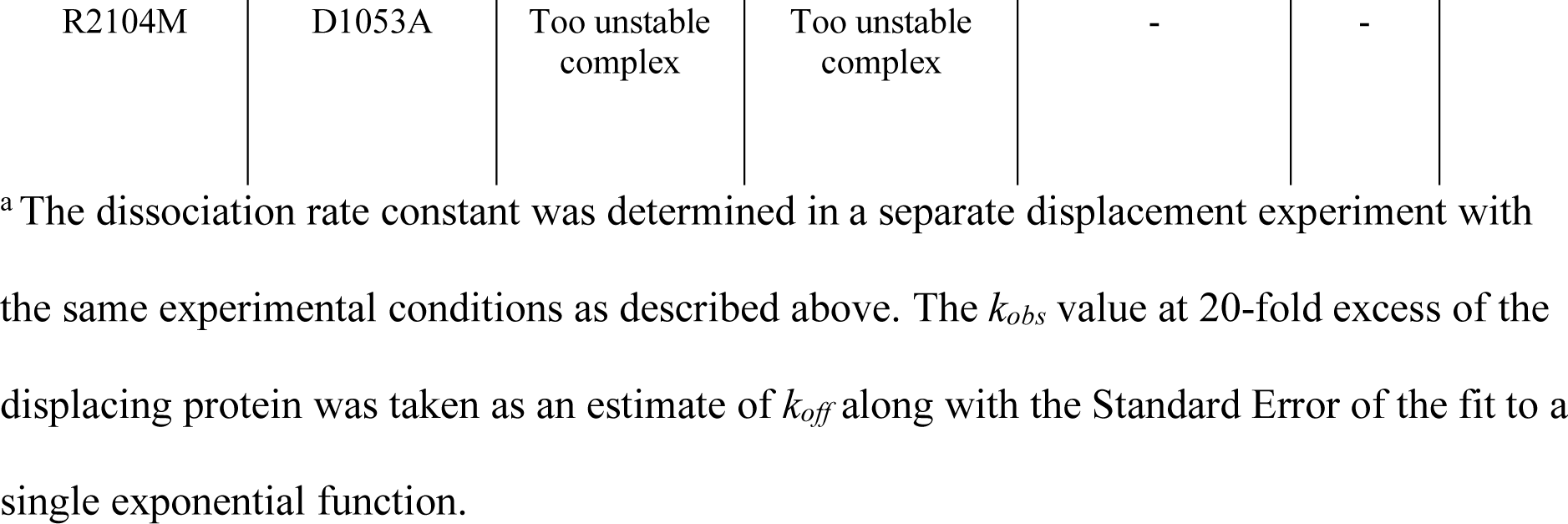
Rate constants of binding for ancestral NCBD and CID variants determined in stopped flow experiments. The mutants were either helix-modulating mutants of ancestral CID or deletion mutations in NCBD or CID chosen to probe for native inter- and intramolecular interactions in the ancestral complex. The rate constants were obtained from global fitting of the kinetic data sets to a simple two-state model by numerical integration and the error is the standard error from the fit. The association rate constant of the wildtype represents an average of five separate experiments and the error is the standard deviation for these measurements. All experiments were performed in 20 mM sodium phosphate pH 7.4, 150 mM NaCl at 4 °C.

| NCBD<sub>D/P</sub> variant | CID<sub>IR</sub> variant | $k_{on}$ ($\mu\text{M}^{-1} \text{s}^{-1}$) | $k_{off}$ ($\text{s}^{-1}$) | Type of mutation | $K_D$ ($\mu\text{M}$) | ϕ-value |
| --- | --- | --- | --- | --- | --- | --- |
| pWT | ML | $30 \pm 3$ | $24.7 \pm 0.3^a$ | - | $0.81 \pm 0.07$ | - |
| pWT | A1047G | $24.8 \pm 0.2$ | $53.8 \pm 0.8$ | Helix modulating Cα1 | $2.17 \pm 0.03$ | $0.21 \pm 0.16$ |
| pWT | D1050A | $31.7 \pm 0.1$ | $21.3 \pm 0.2^a$ | Helix modulating Cα1 | $0.67 \pm 0.01$ | - |
| pWT | D1050G | $18.7 \pm 0.2$ | $72 \pm 0.9$ | Helix modulating Cα1 | $3.85 \pm 0.06$ | $0.30 \pm 0.01$ |
| pWT | S1054A | $39.0 \pm 0.2$ | $14.9 \pm 0.1^a$ | Helix modulating Cα1 | $0.38 \pm 0.003$ | $0.33 \pm 0.21$ |
| pWT | S1054G | $23.2 \pm 0.2$ | $74 \pm 1$ | Helix modulating Cα1 | $3.19 \pm 0.05$ | $0.24 \pm 0.01$ |
| pWT | M1062A | $29.8 \pm 0.2$ | $16.1 \pm 0.3^a$ | Helix modulating Cα2 | $0.54 \pm 0.01$ | $0.05 \pm 0.36$ |
| pWT | M1062G | $26.3 \pm 0.2$ | $21.2 \pm 0.2^a$ | Helix modulating Cα2 | $0.81 \pm 0.01$ | $0.31 \pm 0.06$ |
| pWT | A1065G | $28.5 \pm 0.3$ | $77 \pm 1$ | Helix modulating Cα2 | $2.70 \pm 0.05$ | $0.05 \pm 0.12$ |
| pWT | R1069A | $26.0 \pm 0.1$ | $66.7 \pm 0.6$ | Helix modulating Cα2 | $2.57 \pm 0.03$ | $0.14 \pm 0.13$ |
| pWT | R1069G | 23.0 ± 0.6 | 209 ± 4 | Helix modulating Cα2 | 9.1 ± 0.3 | 0.10 ± 0.04 |
| pWT | L1048A | 13.5 ± 0.1 | 105 ± 1 | Native interactions with Leu2071 | 7.8 ± 0.1 | 0.36 ± 0.07 |
| pWT | L1049A | 18.7 ± 0.1 | 71.8 ± 0.6 | Native interactions with Leu2071, Leu2074 and Phe2100 | 3.84 ± 0.04 | 0.31 ± 0.10 |
| pWT | D1053A | 46.4 ± 0.2 | 22.3 ± 0.2<sup>a</sup> | Electrostatic interaction with Lys2075 and Arg2104 | 0.48 ± 0.005 | 0.81 ± 0.36 |
| pWT | L1055A | 14.0 ± 0.1 | 29.0 ± 0.4<sup>a</sup> | Native interactions with Leu1064 and Leu2074 | 2.07 ± 0.03 | 0.83 ± 0.21 |
| pWT | L1056A | Too unstable complex | Too unstable complex | Native interactions with Leu2074 | - | - |
| pWT | L1064A | Too unstable complex | Too unstable complex | Native interactions with Leu2074 and Phe2100 | - | - |
| pWT | I1067V | 20.4 ± 0.2 | 93 ± 1 | Native interactions with Leu2074, Val2086, Leu2087, Leu2090 and Phe2099 | 4.56 ± 0.07 | 0.23 ± 0.09 |
| pWT | I1067A | Too unstable complex | Too unstable complex | Native interactions with Leu2074, Val2086, Leu2087, Leu2090 and Phe2100 | - | - |
| pWT | D1068A | 39.6 ± 0.2 | 59.5 ± 0.7 | Electrostatic interaction with Arg2104 | 1.50 ± 0.02 | -0.43 ± 0.26 |
| pWT | L1071A | Too unstable complex | Too unstable complex | Native interactions with Leu2071, Leu2074, Val2086, Leu2087, Leu2090, Ala2098, Ala2099 and Ile2101 | - | - |
| L2067A | ML | 14.8 ± 0.2 | 67 ± 1 | Native interactions with Ile2089, Leu2090, Leu2096 | 4.5 ± 0.1 | 0.42 ± 0.10 |
| L2071A | ML | $22.1 \pm 0.1$ | $17.4 \pm 0.1^a$ | Native interactions with Leu2071, Ile2089 and Leu2093 | $0.79 \pm 0.006$ | - |
| L2074A | ML | Too unstable complex | Too unstable complex | Native interactions with Leu2071, Leu2074, Leu2078, Leu2086, Ile2089, Ala2092, Leu2093 and Ile2095 | - |  |
| K2075M | ML | $14.5 \pm 0.1$ | $26.3 \pm 0.4^a$ | Electrostatic interaction with Asp1053 | $1.81 \pm 0.03$ | $0.92 \pm 0.26$ |
| K2075M | D1053A | $18.0 \pm 0.1$ | $26.2 \pm 0.2$ | - | $1.46 \pm 0.01$ | |
| L2086A | ML | Too unstable complex | Too unstable complex | Native interactions with Ile1067, Ala1070 and Leu1071 | - |  |
| L2087A | ML | $15.1 \pm 0.1$ | $26.4 \pm 0.5^a$ | Native interactions with Ala1066, Ile1067, Ala1070 and Leu1071 | $1.75 \pm 0.04$ | $0.91 \pm 0.27$ |
| L2090A | ML | Too unstable complex | Too unstable complex | Native interactions with Ile1067, Ala1070, Leu1071 and Ile1073 | - |  |
| L2096A | ML | $14.9 \pm 0.1$ | $40.7 \pm 0.5$ | Native interactions with Leu2067 and Leu2090 | $2.73 \pm 0.04$ | $0.59 \pm 0.14$ |
| A2099G | ML | $27.6 \pm 0.3$ | $122 \pm 1$ | Native interactions with Leu2067 | $4.42 \pm 0.06$ | $0.06 \pm 0.09$ |
| I2101V | ML | $29.3 \pm 0.1$ | $23.8 \pm 0.3^a$ | Native interactions with Leu1071, Ile1073 and Leu1076 | $0.81 \pm 0.01$ | |
| I2101A | ML | Too unstable complex | Too unstable complex | Native interactions with Leu1071, Ile1073 and Leu1076 | - |  |
| R2104M | ML | Too unstable complex | Too unstable complex | Native interactions with Asp1068 | - |  |

|  |  |  |  |  |  |
| --- | --- | --- | --- | --- | --- |
| R2104M | D1053A | Too unstable complex | Too unstable complex | - | - |

<sup>a</sup> The dissociation rate constant was determined in a separate displacement experiment with the same experimental conditions as described above. The $k_{obs}$ value at 20-fold excess of the displacing protein was taken as an estimate of $k_{off}$ along with the Standard Error of the fit to a single exponential function.

This residue is located in the first *α*-helix, which is transiently populated in the free state of CID and forms many native contacts with NCBD in the transition state^26, 30^. The CID1R in the Cambrian-like complex displayed a similarly high *ϕ*-value (0.83), suggesting that this region forms a conserved nucleus for the coupled binding and folding of CID/NCBD. Further comparison between *ϕ*-values in the Cambrian-like and human complex at a site-by-site basis shows that three mutations in particular, NCBD^L2067A^, NCBD^L2087A^ and NCBD^L2096A^, displayed large differences in *ϕ*-values in the respective complex, with the Cambrian-like complex always showing higher *ϕ*-values. The differences suggest rearrangements of native contacts in the transition states from a low-affinity, more native-like, Cambrian-like complex to a high-affinity, more disordered, human complex (Figure 2 c).

Accordingly, MD simulations showed that the Cambrian-like TS, as compared to the human one, is significantly more compact (as judged by gyration radius, Figure 3e) and less heterogeneous (as judged by pairwise RMSD, Figure 3d), supporting the idea that the TS for formation of the Cambrian-like complex is more native-like. The higher NCBD *ϕ*-values (Table S1 and Figure 2c) measured for the ancestral complex resulted in differences in both the NCBD secondary/tertiary structure and the intermolecular interactions in the TS. In the ancestral TS, the NCBD helix N*α*3 is totally unfolded (consistent with its lower helicity also in the native state)^31^, while helices N*α*1 and N*α*2 are well formed and maintain a native-like relative orientation with numerous N*α*1-N*α*2 contacts (Figure 3c and 3b, box 1). On the other hand, in the human TS, N*α*1 and N*α*2 show lower helical content and contacts are less frequent. The main intermolecular contacts are conserved in human and Cambrian-like TS: in both the cases we observed a stable hydrophobic core (Figure 3b, boxes 2), involving C*α*1 (via residues Leu1052 and Leu1055, which displays high *ϕ*-value in both complexes) and N*α*1 (via residues Leu2071, Leu2074 and Lys2075), but with a slightly different orientation of the two helices. A second relevant interacting region involves the residues of the unstructured CID helices C*α*2-C*α*3, which contact NCBD in both TS ensembles. In the ancestral complex, hydrophobic residues of N*α*2 helix are preferred, while in the human TS the interactions are more dispersed, involving also the longer and structured helix N*α*3 (Figure 3b, box 3). The identified interactions are highly native-like in the case of the ancestral complex (Figure S5) as compared to the human one^24^, where a higher number of transient contacts is observed (Figure S6). These observations agree with a Brønsted plot analysis where *ΔΔ*GTS is plotted versus *ΔΔ*GEQ. A salient feature of a nucleation-condensation mechanism in protein folding, where all non-covalent interactions form cooperatively, is a clear linear dependence of the Brønsted plot. In the case of hydrophobic mutations in Cambrian-like NCBD/CID, the Brønsted plot appeared rather scattered as compared to human NCBD/CID (Fig. 2d) and previously characterized IDP systems. One reason for this could be the relatively narrow range of *ΔΔ*GEQ values that can be obtained for the less stable Cambrian-like complex as compared to human NCBD/CID. Nevertheless, the scatter is consistent with the larger spread of values in the Brønsted plot for Cambrian-like NCBD/CID and a less cooperative disorder-to-order transition of Cambrian-like as compared to human NCBD/CID.

**Figure 3.**
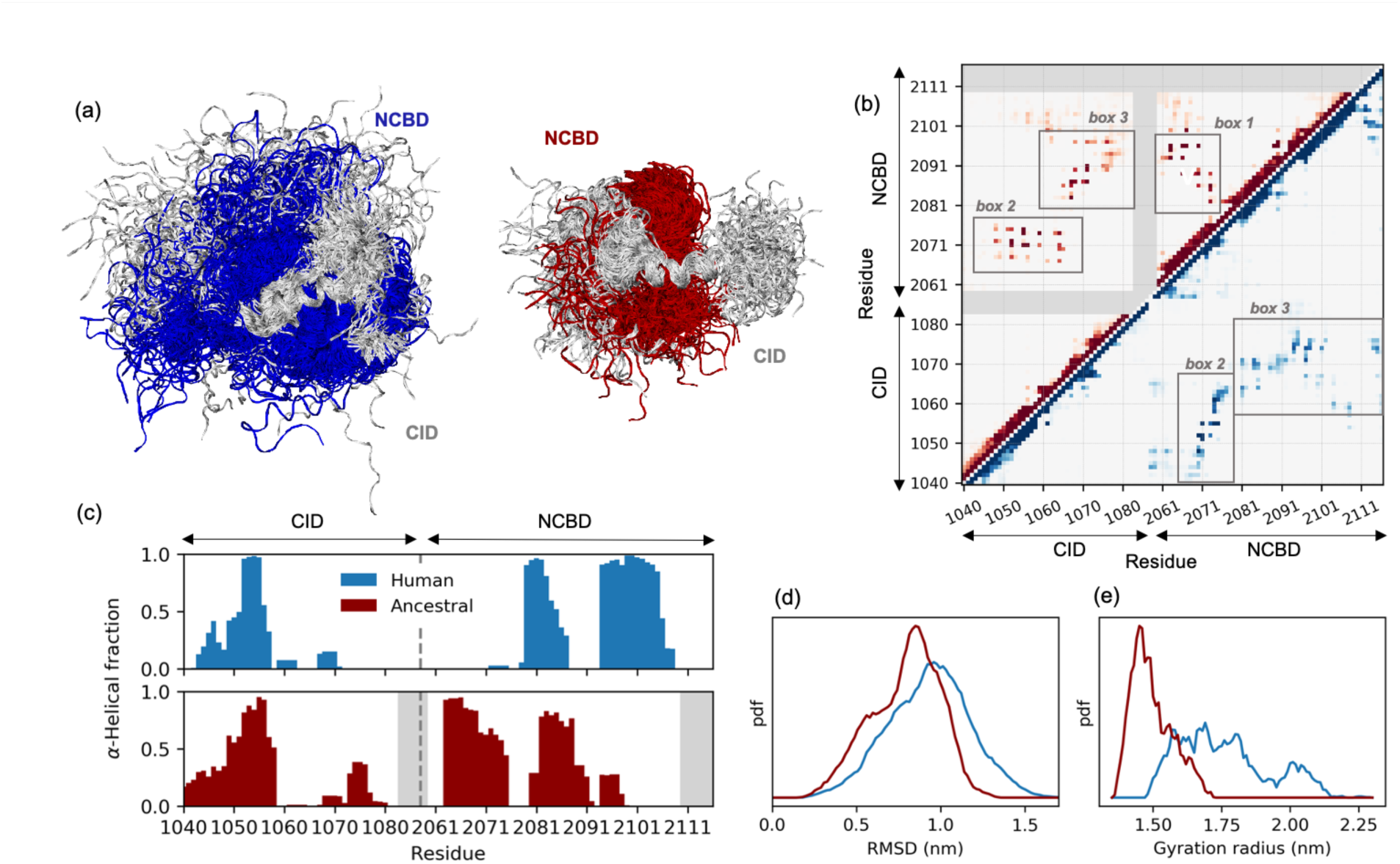
**Comparison of the transition state of human and ancestral Cambrian-like complexes**. (a) MD-determined structural ensembles of the human (NCBD in blue and CID in light grey) and ancestral (NCBD in red and CID in light grey) TS. Both ensembles are aligned on CID helix C*α*1. (b) Map representing the contact probability between each pair of residues in the human (lower right, blue) and in the ancestral (upper left, red) TS ensembles. Probability goes from 0 (white) to 1 (dark blue/red); regions involving residues which are not present in the ancestral complex are shaded with gray. (c) Per-residue *α*-helical content of the human and ancestral TS. (d, e) Probability distribution, in arbitrary units, of the root mean square deviation and of the gyration radius for the human (blue) and ancestral (red) TS ensembles.

### The mechanism of helix formation in CID has been well-conserved during evolution

The binding reaction of NCBD and CID is associated with a dramatic increase in secondary structure content of CID. Helical propensity of the N-terminal helix of CID correlates positively with affinity for NCBD^27^ and modulation of helical propensity is likely an important evolutionary mechanism for tuning affinities of interactions involving IDPs. To investigate native helix formation in the transition state of the Cambrian-like complex, we introduced Ala*→*Gly mutations at surface exposed positions in helix 1 of CID1R (C*α*11R) and in helix 2 (C*α*21R). The mutants were subjected to binding experiments and the rate constants were used to compute *ϕ*-values for each CID variant in complex with NCBDD/P (Table 1). The *ϕ*-values for C*α*11R were ranging between 0.2-0.3 and were lower, close to 0.1, for C*α*21R (Figure 4 a). Brønsted plots resulted in slopes for C*α*11R and C*α*21R of 0.3 and 0, respectively (Figure 4 b). The slope of the Brønsted plot can be regarded as an average *ϕ*-value and thus suggests around 30% and 0% native helical content of C*α*11R and C*α*21R, respectively, in the transition state for the Cambrian-like complex.

**Figure 4.**
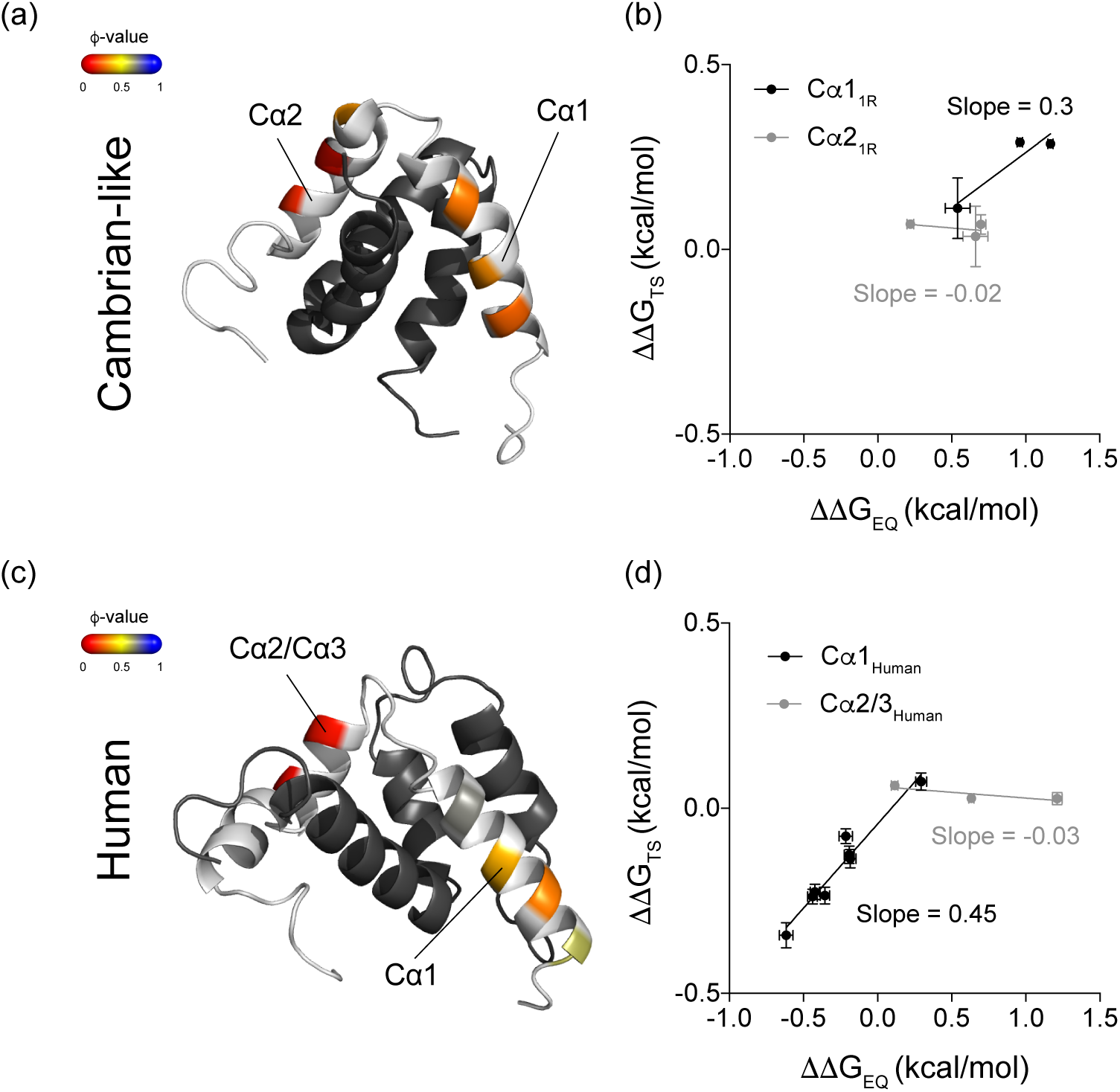
ϕ-values of helix formation in CID in the Cambrian-like and human complex. Ala*→*Gly mutations in surface-exposed positions in the helices of CIDHuman and CID1R were introduced and the kinetic parameters for these helix-modulating mutations were obtained in stopped flow kinetic experiments. The dissociation rate constants were obtained in displacement experiments if *koff* was less than ≈ 30 s^-1^ or otherwise from binding experiments. Using the kinetic parameters (*kon* and *koff*), *ΔΔ*G in the transition state (*ΔΔ*GTS) and in the bound state (*ΔΔ*GEQ) was calculated for each mutant. The experimental conditions were 20 mM sodium phosphate pH 7.4, 150 mM NaCl and the measurements were recorded at 4 °C. (a) *ϕ*-values for helix-modulating mutations in helix 1 (C*α*1) and helix 2 (C*α*2) of CID1R mapped onto the structure of the Cambrian-like protein complex (PDB entry 6ES5)^31^. (b) Brønsted plot for the same helix modulating mutations in CID1R^ML^ in the Cambrian-like complex. The data were fitted with linear regression, yielding slopes of 0.3 ± 0.1 (C*α*1) and −0.02 ± 0.04 (C*α*2). (c) *ϕ*-values for helix-modulating mutations in helix 1 (C*α*1) and helix 2/3 (C*α*2/3) of CID in the human complex mapped onto the structure of the complex (PDB entry 1KBH)^10^. (d) Brønsted plot for helix modulating mutations in helix 1 (C*α*1) and helix 2/3 (C*α*2/3) of human CID in complex with human NCBD. Linear regression analysis yielded slopes of 0.45 ± 0.04 (C*α*1) and −0.03 ± 0.02 (C*α*2/3).

To facilitate a direct comparison with the extant human complex, we extended a previously published data set on helix 1 from human CID (C*α*1Human)^27^ with new Ala*→*Gly mutations in helix 2 (C*α*2Human) and helix 3 (C*α*3Human) (Table 2). The human complex displayed *ϕ*-values for helix formation in C*α*1Human that ranged from 0.3-0.7, whereas all *ϕ*-values in C*α*2Human and C*α*3Human were close to 0 (Figure 4 c). This resulted in Brønsted plots with slopes of 0.5 for C*α*1Human^27^ and virtually 0 for C*α*2Human/C*α*3Human (Figure 4 d). Thus, in both the Cambrian-like and human complexes, the N-terminal C*α*1 plays an important role in forming early intramolecular native secondary structure contacts in the disorder-to-order transition.

These data are well represented by the ancestral TS ensemble. The probability of *α*-helix content in CID, measured via DSSP^32^ and averaged over the whole ensemble, shows that only helix C*α*1 is partially formed in the TS, while other CID helices are mostly unstructured (Figure 3c). Analogously, in the TS of the human complex only helix C*α*1 was folded, overall supporting the importance of C*α*1 formation for NCBD binding.

We note that according to Brønsted plots, C*α*1Human has a slightly higher helical content in the transition state compared to C*α*11R, consistent with a higher helical propensity for C*α*1Human than for C*α*11R, as suggested by predictions using AGADIR^20^. We further note that the A1075G mutation in C*α*3Human did not display a large effect on *KD*, which precluded a reliable estimation of a *ϕ*-value. This could indicate either that C*α*3Human contributes little to the stability of the bound complex or that the Ala*→*Gly substitution at this position promotes an alternative conformation that binds with equal affinity as the wildtype protein. Similarly, Val1077*→*Ala in C*α*3Human was shown previously to have a small positive effect on the affinity for NCBD^29^, suggesting structural re-arrangement in the bound state.

### Role of a conserved and buried salt-bridge in the Cambrian-like complex

Long-range electrostatic interactions promote association of proteins and play a major role in IDPs. Mutation of a conserved salt-bridge between Arg2104 in NCBD and Asp1068 in CID was previously shown to display large effects on the kinetics of complex formation for human NCBD/CID, both in terms of a 10-fold reduction in *k*on but also with the occurrence of a new kinetic phase (*kobs* ≈ 15-20 s^-1^). Analysis of the kinetic data favored an induced fit model, thus a conformational change after binding^16, 33, 34^. We generated the protein variants NCBDDR2104M _and CID_ ^D1068A^ to assess the role of this salt-bridge in the ancestral/P 1R Cambrian-like complex.

NCBDD/P and CID1R displayed clear biphasic kinetic traces in the stopped flow experiments, similarly to experiments with the corresponding mutants in the human complex. Fitting of the kinetic data to obtain *kobs* values revealed one concentration-dependent kinetic phase, which increased linearly with CID concentration and a second kinetic phase which was constant at *kobs* ≈ 16 s^-1^ over the entire concentration range. Thus, the kinetic data set for the complex between NCBDD/P^R2104M^/CID1R was fitted globally to an induced fit mechanism to obtain estimates of the microscopic rate constants. The comparison between the Cambrian-like and human complex revealed that the effect of mutating the buried salt-bridge was much smaller with regard to the association rate constant *k1* for the Cambrian-like complex than for the human complex, less than 2-fold versus 20-fold, respectively. On the other hand, the slow phase was similar for both the human and ancestral complex with a *k*obs (= *k2* + *k-2*) of 15-20 s^-1^. However, global fitting suggested that the alternative conformation of the bound state is only slightly populated for the Cambrian-like complex (*k-2*>>*k2)*. This was corroborated by ITC measurements, which showed that the overall *KD* (5.1 ± 0.3 µM) is highly consistent with *k-1*/*k1* (4.8 µM). Interestingly, while the salt-bridge is significantly populated in the native state simulations, it is not populated in neither the Cambrian-like nor the human TS ensemble, suggesting that the formation of this interaction is not relevant for the initial recognition. Nonetheless these residues promote the association of the human complex most likely via unspecific long-range interactions.

Furthermore, we mutated Asp1053 in CID ^ML^ to Ala, to assess potential salt-bridge formation between this residue and Arg2104 in NCBDD/P. The complex between NCBDD/P and CID1R was very destabilized and the kinetic phase that reported on the binding event was too fast for the stopped-flow instrument. However, the concentration-independent kinetic phase was detected (*kobs* ≈ 20-40 s^-1^). Single charge mutations cannot be considered conservative since they may result in unpaired charges in or close to hydrophobic interfaces. Any effects from such mutations may also be due to non-specific charge-charge attraction or repulsion. (The overall charge of NCBDD/P is positive and CID1R is negative.) Nevertheless, we report kinetic data for such single mutants. Interestingly, CID_IR_^D1068A^ displayed a negative *ϕ*-value (ca. −0.4, Table 1 and Table 3) due to an increase in both *k*on and *k*off upon mutation, suggesting that Asp1068 makes a non-favorable interaction in the transition state. This is consistent with the small effect on *k*on for NCBDD/P /CID1R, which might result from opposing effects on *k*on by the respective mutation. The other Asp mutant, CID1R, also displayed an increase in *k*on but a positive high *ϕ*-value. Thus, Asp1053 forms non-favorable interactions both in the transition state and in the native state of the complex. The NCBDD/P variant gives a high *ϕ*-value suggesting a native interaction in the transition state. While Asp1053 is in the vicinity of Lys2075 the coupling free energy between them is low (0.17 kcal/mol) and it is not clear what interactions these residues make in the native complex. All surface charge mutations had little effect on kinetics or yielded low *ϕ*-values suggesting that overall charge plays a minor role in the association of NCBDD/P and CID1R (Table 1) and similarly for human NCBD/CID (Table 2).

**Table 2.**
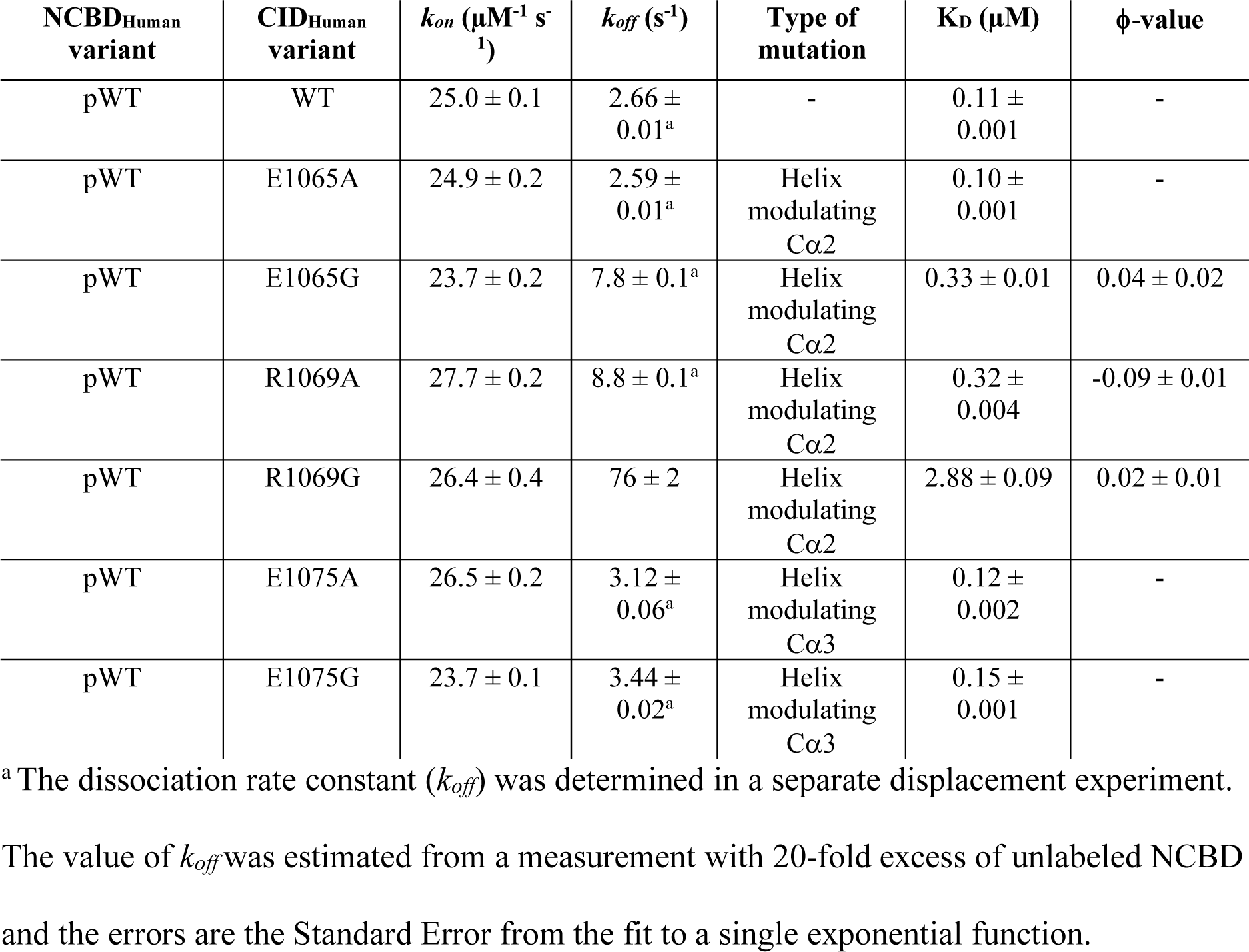
Rate constants for helix-modulating mutations in helix 2 and helix 3 of human CID. The rate constants were obtained in fluorescence-monitored stopped flow kinetic experiments. The experimental conditions were 20 mM sodium phosphate buffer pH 7.4, 150 mM NaCl and the experiments were performed at 4°C. The errors are Standard Errors from global fitting to a simple two-state model.

**Table 3.** Rate constants of binding of NCBDD/P to different CID variants determined in stopped flow experiments. The errors are Standard Errors from global fitting of the kinetic data to a two-state model. All experiments were performed in 20 mM sodium phosphate pH 7.4, 150 mM NaCl at 4 °C. In general, the resulting *ϕ*-values are similar to the ones obtained using the NCBDD/P^pWT^ variant with Trp2073 (shown in parenthesis).

| NCBD<sub>D/P</sub><sup>pWT</sup><br>variant | CID<sub>IR</sub><br>variant | $k_{on}$ ($\mu\text{M}^{-1} \text{s}^{-1}$) | $k_{off}$ ($\text{s}^{-1}$) | Type of<br>mutation | $K_D$ ($\mu\text{M}$) | ϕ-value |
| --- | --- | --- | --- | --- | --- | --- |
| L2067W | ML | $17.9 \pm 0.1$ | $39.2 \pm 0.2$ | - | $2.2 \pm 0.02$ | - |
| L2067W | A1047G | $12.9 \pm 0.1$ | $64.7 \pm 0.6$ | Helix<br>modulating<br>Cα1 | $5.0 \pm 0.1$ | $0.40 \pm 0.02$<br>($0.21 \pm 0.16$) |
| L2067W | L1049A | $9.6 \pm 0.1$ | $52.9 \pm 0.2$ | Native<br>interactions<br>with<br>Leu2071,<br>Leu2074 and<br>Phe2100 | $5.5 \pm 0.1$ | $0.68 \pm 0.03$<br>($0.31 \pm 0.17$) |
| L2067W | D1053A | $29.3 \pm 0.1$ | $25.7 \pm 0.1$ | Electrostatic<br>interaction<br>with Lys2075<br>and Arg2104 | $0.88 \pm 0.01$ | $0.54 \pm 0.02$<br>($0.81 \pm 0.36$) |
| L2067W | I1067V | $10.7 \pm 0.1$ | $132 \pm 1$ | Native<br>interactions<br>with<br>Leu2074,<br>Val2086,<br>Leu2087,<br>Leu2090 and<br>Phe2099 | $12.3 \pm 0.1$ | $0.30 \pm 0.01$<br>($0.23 \pm 0.09$) |
| L2067W | A1065G | $14.4 \pm 0.2$ | $125 \pm 1$ | Helix<br>modulating<br>Cα2 | $8.7 \pm 0.1$ | $0.16 \pm 0.02$<br>($0.05 \pm 0.12$) |
| L2067W | D1068A | $27.0 \pm 0.3$ | $178 \pm 2$ | Electrostatic<br>interaction<br>with Arg2104 | $6.6 \pm 0.1$ | $-0.37 \pm 0.02$<br>($-0.43 \pm 0.26$) |

In agreement with these data, simulations supported the idea that hydrophobic interactions are more relevant than electrostatic contacts in the TS for formation of the ancestral complex. In fact, no stable salt-bridges or hydrogen bonds were observed, with Arg2104 contacting different polar residues only in a transient manner. Also, the high *ϕ*-value of the NCBDD/P variant can be explained by the ability of Lys2075 to engage in hydrophobic, rather than polar, interactions stabilizing the native-like hydrophobic core formed by helices C*α*1-N*α*1 (Figures 3b and S5).

**Figure 5.**
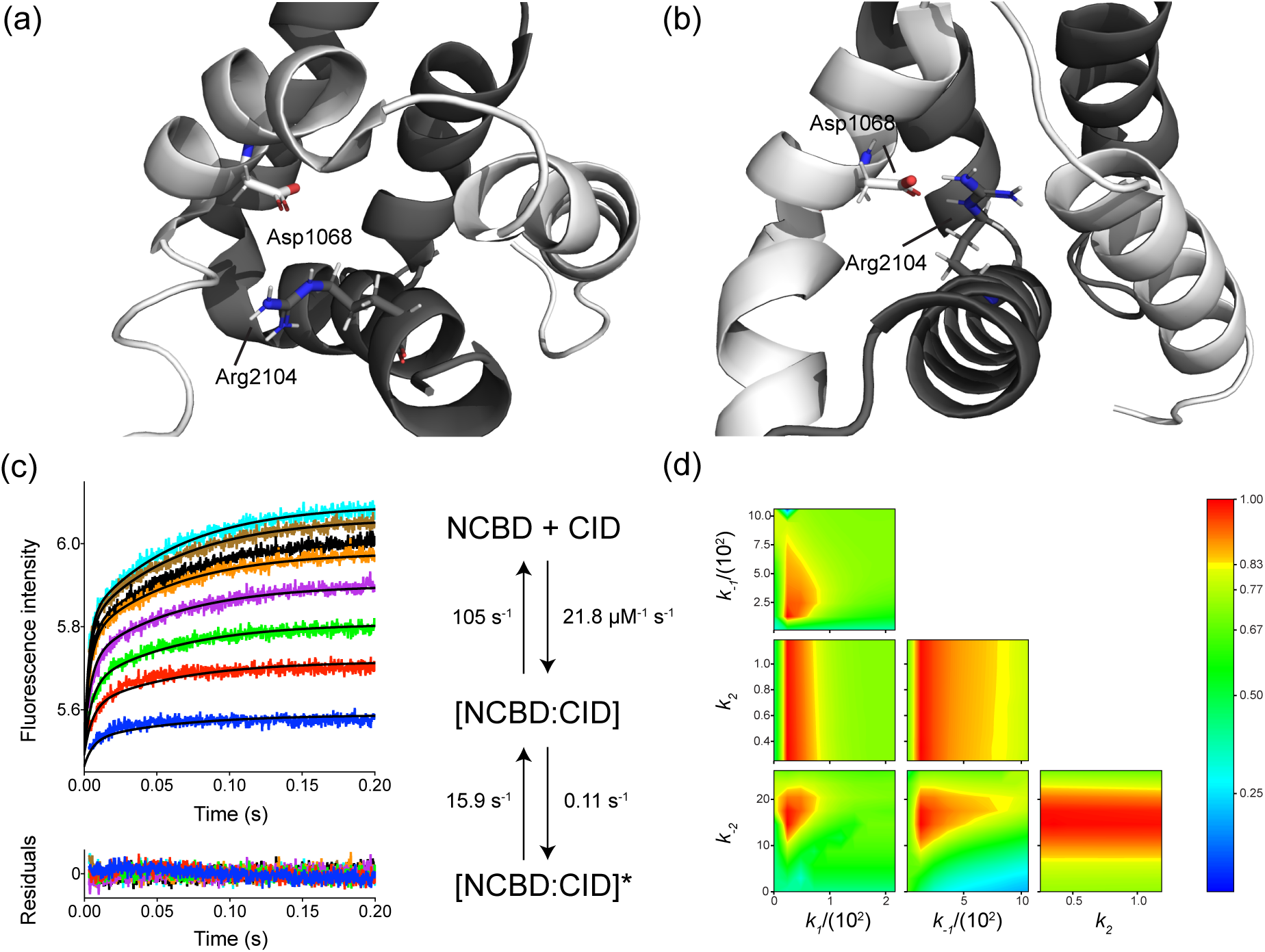
The conserved salt-bridge between Arg2104 in NCBD and Asp1068 in CID. The structure of (a) the Cambrian-like (PDB entry 6ES5)^21^ and (b) the human NCBD/CID complex with Arg2104 and Asp1068 forming the salt-bridge highlighted as stick model. NCBD is in dark grey and CID in light grey. (c) The stopped-flow experiment for the NCBDD/P /CID1R complex was performed in 20 mM NaPi pH 7.4, 150 mM NaCl at 4° C. The stopped flow kinetic traces were fitted globally to an induced fit model in order to obtain microscopic rate constants for each reaction step. The black solid lines represent the best fit to the kinetic traces and the residuals are shown below the curve. The best-fit microscopic rate constants obtained from global fitting to an induced fit model are shown next to the binding curves. (d) The confidence contour plot shows the variation in chi^2^ as two parameters are systematically varied while the rest of the parameters are allowed to float, which can reveal patterns of co-variation between parameters in a model. The color denotes the chi^2^/chi^2^min value according to the scale bar to the right. Here, the confidence contour plot showed that *k2* was poorly defined. The yellow boundary represents a cutoff in chi^2^/chi^2^min of 0.8.

## Discussion

The higher prevalence of IDPs among eukaryotes as compared to prokaryotes suggests that these proteins have played an important role in the evolution of complex multicellular organisms^35, 36^. IDPs often participate in regulatory functions in the cell, by engaging in complex interaction networks that fine-tune cellular responses to environmental cues^37^. One feature common to many IDPs, and which has likely contributed to their abundance in regulatory functions, is the ability to interact specifically with several partners that are competing for binding^38^. NCBD is an archetype example of such a disordered protein interaction domain that has evolved to bind several cellular targets, including transcription factors and transcriptional co-regulators^10, 11, 39^. Every time a new partner was included in the repertoire, NCBD somehow adapted its affinity for the new ligand, while maintaining affinity for already established one(s), as occurred around 450-500 Myr, when the interaction between NCBD and CID was established^20^. On a molecular level, it is intriguing how such multi partner protein domains evolve. In the present study, we have extended our structural studies^31^ and investigated the evolution of the binding mechanism using site-directed mutagenesis, *ϕ*-value analysis and restrained MD simulations to shed light on changes occurring at the molecular level when the low-affinity Cambrian-like NCBD evolved higher affinity for its protein ligand CID.

Recent works suggest that IDPs can adopt multiple strategies for recognizing their partners. Gianni and co-workers proposed the concept of templated folding, where the folding of the IDP is modulated, or templated, by its binding partner, as shown for cMyb/KIX^40^, MLL/KIX^41^ and NTAIL/XD^42, 43^. Similar ideas were put forward by Zhou and coworkers based on experiments on WASP GBD/Cdc42 and formulated in terms of multiple dock-and-coalesce pathways^44^. On the other hand, studies on disordered domains from BH3-only proteins binding to BCL-2 family proteins suggest conservation of *ϕ* values and a more robust folding mechanism^45^. We recently showed by double mutants and simulation that a high plasticity in terms of formation of native hydrophobic interactions in the transition state exists for human NCBD/CID^15^ where both partner are very flexible, in agreement with templated folding.

In the present study, by comparing the TSs for formation of human and Cambrian-like NCBD/CID complexes, we demonstrate that, while similar core interacting regions have been conserved throughout evolution, the interaction between the two proteins has evolved from a more ordered ancestral TS to the heterogenous and plastic behavior observed in the human complex. We find that the fraction of CID helical content in the transition state is overall conserved, with intermediate values in C*α*1 (slightly higher in human than in Cambrian-like NCBD/CID) and low values in C*α*2/3. Conversely, we observe that the transition state of the low-affinity Cambrian-like complex has more native-like features in terms of hydrophobic interactions (higher *ϕ*-values) as compared to the human one, with clear site-specific differences such as residues Leu2067 Leu2087 and Leu2096 of NCBD. In the ancestral TS, fewer but more native-like contacts are required to be formed (Figure S5 and S6) and proper NCBD tertiary structure (regulating N*α*1-N*α*2 orientation) is achieved before CID binding. Vice-versa, in human TS numerous transient inter-molecular interactions are engaged (Figure S6), involving a large number of residues of both CID and NCBD.

In the homeodomain family of proteins a spectrum of folding mechanisms ranging from nucleation-condensation to diffusion-collision was previously observed^46^. Furthermore, it has been suggested that the two mechanism can be related to the balance between hydrophobic and electrostatic interactions^47^. Mutational studies on IDPs^13, 40, 41, 48–53^ are more or less consistent with apparent two-state kinetics and the nucleation-condensation mechanism of globular proteins^54^, i.e. cooperative, simultaneous formation of all non-covalent interactions around one well defined core. Salient features of this mechanism are linear Brønsted plots and fractional *ϕ*-values. One alternative mechanism would be independently folding structural elements, which dock to form the tertiary structure as formulated in the diffusion-collision model^46, 55^. In such scenario, Brønsted plots would be more scattered and the *ϕ*-values be both low and high and clustered in structurally contiguous contexts and even separated into two or more folding nuclei. Our comparison of secondary and tertiary structure formation in human versus Cambrian-like NCBD/CID is therefore interesting since it shows that the folding of certain elements of secondary structure can be distinct from others, and that they may or may not be part of an extended folding nucleus. The Brønsted plot for Cambrian-like NCBD/CID shows a larger scatter than that for human NCBD/CID (Fig. 2 d). Whereas C*α*1 of both CID1R and human CID displays fractional *ϕ*-values, C*α*2/3 have *ϕ*-values of zero (Fig. 4).

Thus, C*α*1 may function as a well-defined folding nucleus around which remaining structure condensate, as observed for high-affinity human NCBD. However, C*α*1 may also be part of a more extended folding nucleus together with hydrophobic tertiary interactions as in low-affinity Cambrian-like NCBD/CID (Fig. 2-4), but not to the extent that we define it as two separate folding nuclei. The Cambrian-like NCBD/CID shows therefore an intermediate behavior between nucleation-condensation and diffusion-collision mechanism, that shifted towards nucleation-condensation during evolution. Three other IDP interactions with more than one helical segment, Hif-1*α* CAD^53^, TAD-STAT2^52^, and pKID^56^ (all binding to KIX), do not display this behavior, but show mainly low *ϕ*-values (<0.2) with only one or a few higher ones. Thus, so far, nucleation-condensation appears more prevalent for globular protein domains^57^, as well as for IDPs in disorder-to-order transitions. It will be interesting to see whether other IDPs with several secondary structure elements display any distinct distribution of *ϕ*-values.

## Materials and methods

### Ancestral and human protein sequences

The reconstruction of ancestral sequences of NCBD (from the CREBBP/p300 protein family) and CID (from the NCOA/p160/SRC protein family) has been described in detail before^20^. Briefly, protein sequences in these families from various phyla were aligned and the ancestral protein sequences of NCBD and CID were predicted using a maximum likelihood (ML) method. The ML ancestral protein variants of NCBD and CID, NCBDD/P and CID1R, were used as “wildtypes” in this study. The human NCBD protein was composed of residues 2058-2116 from human CREBBP (UniProt ID: Q92793) and the human CID protein was composed of residues 1018-1088 from human NCOA3/ACTR (UniProt ID: Q9Y6Q9), in accordance with previous studies on the human protein domains^16, 26, 29^. The reconstructed ancestral sequences of NCBD and CID were shortened to contain only the evolutionarily more well-conserved regions that form a well-defined structure upon association with the other domain. Thus, the ancestral NCBD variant was composed of residues corresponding to 2062-2109 in human CREBBP and the ancestral CID variant was composed of residues corresponding to 1040-1081 in human NCOA3/ACTR.

### Cloning and mutagenesis

The cDNA sequences for the protein variants used in the study were purchased from GenScript and the proteins were N-terminally tagged with a 6xHis-Lipo domain. The mutants were generated using a whole plasmid PCR method. The primers were typically two complementary 33-mer oligonucleotides with mis-matching bases at the site of the mutation, which were flanked on each side by 15 complementary bases. The annealing temperature in the PCR reactions was between 55-65 °C and the reactions were run for 20 cycles. The products were transformed into *E. coli* XL-1 Blue Competent Cells and selected on LB agar plates with 100 µg/mL ampicillin. The plasmids were purified using the PureYield™ Plasmid Miniprep System (Promega).

### Protein expression and purification

The plasmids encoding the protein constructs were transformed into *E. coli* BL-21 DE3 pLysS (Invitrogen) and selected on LB agar plates with 35 µg/mL chloramphenicol and 100 µg/mL ampicillin. Colonies were used to inoculate LB media with 50 µg/mL ampicillin and the cultures were grown at 37 °C to reach OD600 0.6-0.7 prior to induction with 1 mM isopropyl *β*-D-1-thiogalactopyranoside and overnight expression at 18 °C. The cells were lysed by sonication and centrifuged at approximately 50,000 *g* to remove cell debris. The lysate was separated on a Ni Sepharose 6 Fast Flow (GE Healthcare) column using 30 mM Tris-HCl pH 8.0, 500 mM NaCl as the binding buffer and 30 mM Tris-HCl pH 8.0, 500 mM NaCl, 250 mM imidazol as the elution buffer. The 6xHis-Lipo tag was cleaved off using Thrombin (GE Healthcare) and the protein was separated from the cleaved tag using the same column and buffers as described above. Lastly, the protein was separated on a RESOURCE™ reversed phase chromatography column (GE Healthcare) using a 0-70 % acetonitrile gradient. The purity of the protein was verified by the single-peak appearance on the chromatogram or by SDS-PAGE. The identity was verified by MALDI-TOF mass spectrometry. The fractions containing pure protein were lyophilized and the concentration of the protein was measured by absorption spectrometry at 280 nm for variants that contained a Tyr or Trp residue. For the proteins which lacked a Tyr or Trp residue, absorption at 205 nm was used to estimate the concentration. The extinction coefficient for human CID was previously determined by amino acid analysis to 250,000 M^-1^ cm^-1^ at 205 nm. For the shorter ancestral variants, the extinction coefficient was calculated based on the amino acid sequence^58^.

### Design and evaluation of NCBDD/P Trp variants

In order to perform fluorescence-monitored stopped-flow kinetic experiments, a fluorescent probe is required. As both NCBD and CID lack Trp residues, which provides the best sensitivity in fluorescence-monitored experiments, several NCBDD/P^ML^ variants with Trp residues introduced at different positions were constructed: NCBDD/P^L2067W^, NCBDD/P, NCBDD/P, NCBDD/P and NCBDD/P. The NCBDD/P variant corresponds to the NCBDHuman^Y2108W^ variant, which was used previously as a “pseudo-wildtype” in stopped flow kinetic experiments^15, 16, 29^. These NCBDD/P^ML^ Trp variants were assessed based on secondary structure content and stability of complex with CID using far-UV circular dichroism (CD) spectroscopy, and by kinetic and equilibrium parameters from stopped-flow fluorescence spectroscopy and isothermal titration calorimetry (ITC), in order to find an engineered NCBDD/P^ML^ variant with similar biophysical properties to the wildtype NCBDD/P (Figure S1). The NCBDD/P variant displayed the most similar behavior to NCBDD/P. Our data showed that the structural content, complex stability as well as affinity of this Trp variant was highly similar to NCBDD/P^ML^ (Figure S1). Thus, our data validated the use of NCBDD/P variant as a representative of NCBDD/P and all stopped flow kinetic experiments for the ancestral Cambrian-like complex were performed using the “pseudo-wildtype” NCBDD/P^T2073W^ variant, which we denote NCBDD/P.

### Stopped-flow spectroscopy and calculation of ϕ-values

The kinetic experiments were conducted using an upgraded SX-17MV Stopped-flow spectrofluorometer (Applied Photophysics). The excitation wavelength was set to 280 nm and the emitted light was detected after passing through a 320 nm long-pass filter. All experiments were performed at 4 °C and the default buffer for all experiments was 20 mM sodium phosphate pH 7.4, 150 mM NaCl. In order to promote secondary and tertiary structure formation of some structurally destabilized NCBD mutants, the experimental buffer was supplemented with 0.7 M trimethylamine N-oxide (TMAO; Figure S2 c-d). Typically, in kinetic experiments the concentration of NCBD was kept constant at 1-2 µM and the concentration of CID was varied between 1-10 µM. Experiments where the concentration of CID was kept constant while NCBD was varied were also performed to check for consistency of the obtained results. In these experiments, CID was held constant at 2 µM and NCBD was varied between 2-10 µM. The kinetic binding curves with NCBD in excess was in good agreement with those using CID in excess, but since the quality of the kinetic traces were better when CID was in excess, these experiments were used to determine the kinetic parameters for the different variants reported in the paper. All experiments were performed using the pseudo wildtype NCBD variants, which was the NCBDD/P and NCBDHuman^Y2108W^ variants. These variants are denoted as NCBDD/P and NCBDHuman, respectively.

The dissociation rate constant *koff* can be determined from binding experiments, but the accuracy decreases whenever *kobs* values are much larger than *koff* or when *koff* is very low. In the present study, displacement experiments were performed for protein complexes with *koff* values below 30 s^-1^ in binding experiments. In displacement experiments, an unlabeled NCBD variant (without Trp) was used to displace the different NCBDD/P^pWT^ or NCBDHuman^pWT^ variants from the complexes. The *kobs* value at 20-fold excess of the unlabeled NCBD variant was taken as an estimate of the dissociation rate constant, *koff*.

The *ϕ*-values for each mutant were computed using the rate constants that were obtained in the stopped-flow measurements (e.g *kon* and *koff*) using Equations 1-3.

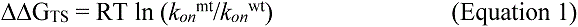

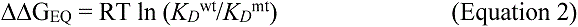

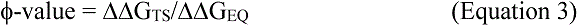

### CD spectroscopy

Far-UV circular dichroism (CD) spectra were acquired using a J-1500 spectrophotometer (JASCO) in 20 mM sodium phosphate buffer pH 7.4, 150 mM NaCl at 4 °C. The bandwidth was 1 nm, scanning speed 50 nm/min and data pitch 1 nm. The protein concentrations were between 20-40 µM for all protein variants and each spectrum was typically an average of 2-3 individual spectra. The thermal denaturation experiments of the protein complexes were performed by monitoring the CD signal of 20 µM NCBD in complex with 20 µM CID at 222 nm in the same experimental buffer as above and over a temperature range of 4-95 °C. For these experiments, the heating speed was 1 °C/min with 5 seconds waiting time at each data point and data was acquired every 1 °C.

### NMR spectroscopy

NMR samples were prepared by mixing the labelled NCBD or CID solution with excess amount of unlabeled CID or NCBD solution, followed by lyophilization and rehydration. The final samples had a labelled NCBD or CID concentration of approximately 0.5 mM and a phosphate buffer concentration of 20 mM at pH 7. During dissolution, 0.01% NaN3 and 10% D2O were added. All NMR spectra were recorded at 25 °C on a 600 MHz Bruker Avance Neo NMR spectrometer equipped with a TCI cryo-probe. The ^1^H ^15^N HSQC spectra were collected with 2048 data points in *ω*2 /^1^H dimension and 256 data points in *ω*1/^15^N dimension; 4 or 8 scans were taken. All spectra were processed with TopSpin 3.2 and analyzed with Sparky 3.115^59^. During this analysis, a downfield shift of 1.0 ppm in ^15^N dimension and a downfield shift of 0.13 ppm in ^1^H dimension were specifically applied to the ppm scale for ^15^N HSQC spectrum of unlabeled CID1R bound to N-labelled NCBDD/P.

### Isothermal titration calorimetry

Isothermal titration calorimetry measurements were performed at 25 °C in a MicroCal iTC200 System (GE Healthcare). The proteins were dialyzed simultaneously in the same experimental buffer (20 mM sodium phosphate pH 7.4, 150 mM NaCl) in order to reduce buffer mismatch. The concentration of NCBD in the cell was 12-50 µM (depending on variant) and the concentration of CID in the syringe was between 120-500 µM, depending on NCBD concentration, such that a 1:2 stoichiometry was achieved at the end of each experiment. The data were fitted using the built-in software to a two-state binding model.

### Data analysis using numerical integration

The stopped flow kinetic data sets were fitted using the KinTek Explorer software (KinTek Corporation)^60, 61^. The software employs numerical integration to simulate and fit reaction profiles directly to a mechanistic model. Scaling factors were used to correct for small fluctuations in lamp intensity and errors in concentration, but they were generally close to 1. In cases where the signal-to-noise in the obtained data was low, scaling factors were not applied. For some more-than-one-step models, two-dimensional confidence contour plots were computed to assess confidence limits for each parameter and co-variation between parameters. An estimated real time zero of the stopped flow instrument of −1.25 ms was used to adjust the timeline in order to obtain correct kinetic amplitudes. The fitted data was exported and graphs were created in GraphPad Prism vs. 6.0 (GraphPad Software).

### MD simulations of the transition state ensembles

The transition state for formation of the human complex was previously determined by means of *ϕ*-value restrained molecular dynamics simulations^24^. Here, the same procedure was followed to determine the TS of the ancestral CID-NCBD complex. The simulations were performed with GROMACS 2018^62^ and the PLUMED2 software^63^, using the Amber03w force field^64^ and the TIP4P/2005 water model^65^. The initial conformation was taken from available PDB structure (6ES5)^31^ and modified with Pymol^66^ to account for the T2073W mutation. The structure was solvated with ∼6700/16800 water molecules (for native state and TS simulations, respectively), neutralized, minimized and equilibrated at the temperature of 278 K using the Berendsen thermostat^67^. Production simulations were run in the canonical ensemble, thermosetting the system using the Bussi thermostat^68^; bonds involving hydrogens were constrained with the LINCS algorithm^69^, electrostatic was treated by using the particle mesh Ewald scheme^70^ with a short-range cut-off of 0.9 nm and van der Waals interaction cut-off was set to 0.9 nm.

A reference native state simulation, at the temperature of 278 K, was performed to determine native contacts. Firstly, we ran a 40 ns long restrained simulation to enforce agreement with atomic inter-molecular upper distances previously determined from NMR experiments^31^: to this aim lower wall restraints were applied on the NOE-converted distances. Subsequently, an unrestrained 280 ns long simulation was performed and the last 200 ns were used to determine native contacts: given two residues that are not nearest neighbors, native contacts are defined as the number of heavy side-chain atoms within 0.6 nm in at least 50% of the frames.

The TS ensemble of the ancestral complex was determined via *ϕ*-value restrained MD simulations, following a standard procedure based on the interpretation of *ϕ*-value analysis in terms of fraction of native contacts^24, 71, 72^. Herein, restraints (in the form of a pseudo energy term accounting for the square distance between experimental and simulated *ϕ*-values) are added to the force field to maximize the agreement with the experimental data: the underlying hypothesis is that structures reproducing all the measured *ϕ*-values are good representations of the TS. From each conformation the *ϕ*-value for a residue is back-calculated as the fraction of the native contact (determined from the native state simulation) that it makes, implying that only *ϕ*-values between 0 and 1 can be used as restraints. Totally, we included 11 *ϕ*-values in this range, all based on single conservative point mutations. Mutations involving charged amino acids (namely, K2075M, involved in intermolecular interactions, and the Ala*→*Gly substitutions at positions D1050 and R1069, probing the helical content of CID helices C*α*1 and C*α*2, respectively) were excluded; we however verified that the structural ensemble obtained could provide a consistent interpretation of the associated *ϕ*-values. A list of the *ϕ*-values used in the ancestral and human TS simulations is reported in Table S1. The TS ensemble was generated using simulated annealing, performing 1334 annealing cycles, each 150 ps long, in which the temperature was varied between 278 K and 378 K, for a total simulation time of 200 ns. The TS was determined using only the structures sampled at the reference temperature of 278 K in the last 150 ns of simulation, resulting in an ensemble of ∼5400 conformations.

## Acknowledgements

This work was funded by the Swedish Research Council grant 2016-04965 (to P.J.). We used the NMR Uppsala infrastructure, which is funded by the Department of Chemistry - BMC and the Disciplinary Domain of Medicine and Pharmacy. C.C. acknowledges CINECA for an award under the ISCRA initiative, for the availability of high-performance computing resources and support.

## Author contributions

E.K. and P.J. conceived and designed the project. E.K., A.E., Z.A.T., F.S., E.A. and W.Y. performed experiments and analyzed data. C.P. and C.C. designed, performed and analyzed all MD simulations. E.K., C.P., C.C., and P.J. interpreted the data and wrote the paper.

## Competing interests

The authors declare no competing interests.

## Supplementary information

**Figure S1.**
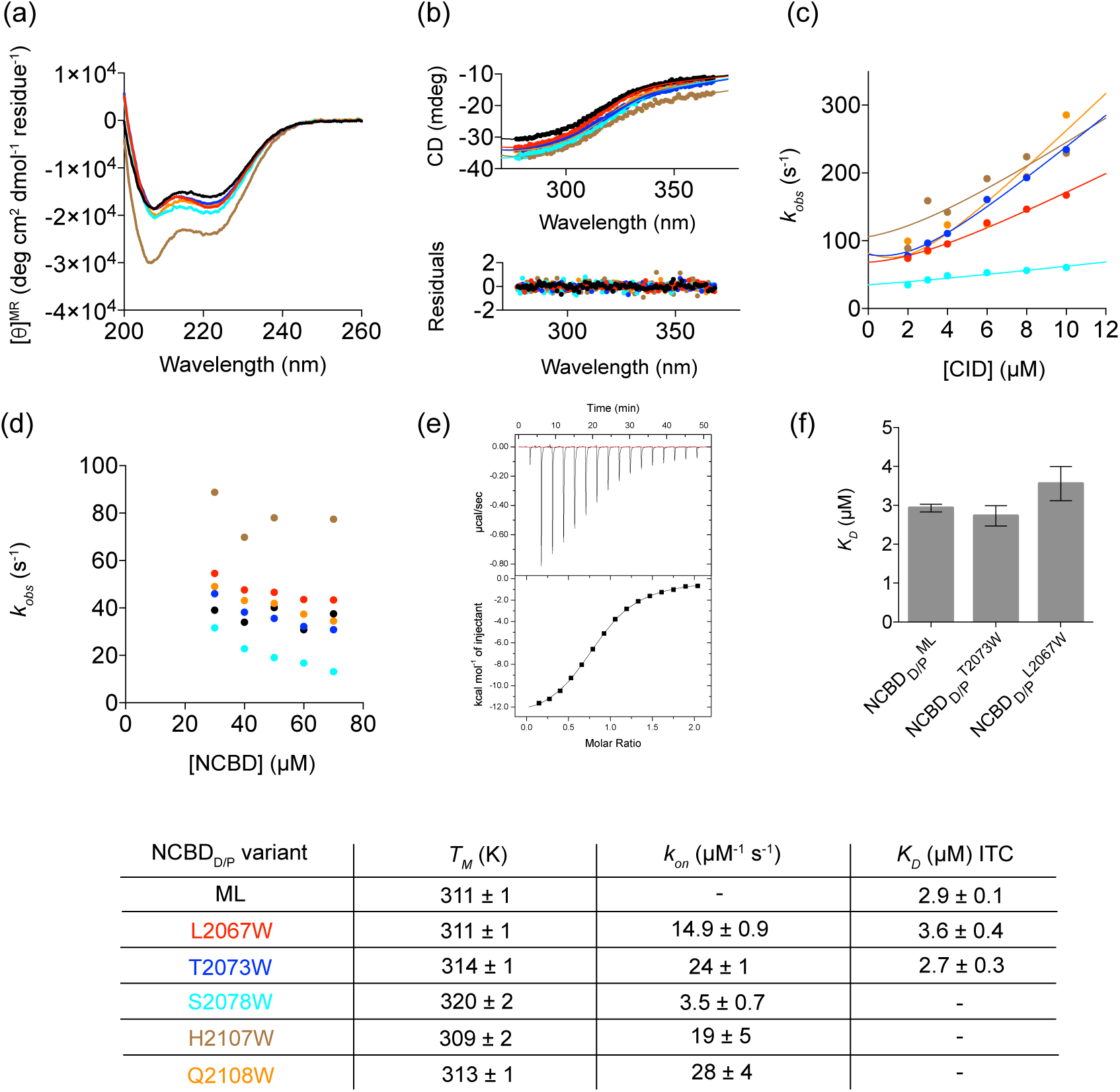
Structural, thermodynamic and kinetic properties of different NCBDD/P Trp variants. For all datasets, the color coding is the following: NCBDD/P^ML^ (black), NCBDD/P (red), NCBDD/P (blue), NCBDD/P (cyan), NCBDD/P (brown) and NCBDD/P^Q2108W^ (orange). All experiments were conducted in 20 mM sodium phosphate pH 7.4, 150 mM NaCl at 4°C unless otherwise stated. (a) CD spectra for NCBDD/P and the different NCBDD/P Trp variants. (b) Thermal stability of different NCBDD/P Trp variants in complex with CID1R. The CD signal at 222 nm was used to monitor complex dissociation and the temperature interval was 4-95 °C. The data were fitted to a two-state model (solid line, residuals in lower panel) in order to obtain estimates of the thermal denaturation midpoint of the respective NCBD/CID complex. (c) Observed rate constants (*k*obs) from stopped flow binding experiments for NCBDD/P Trp variants and CID1R. The concentration of NCBD was held constant at 2 µM and the concentration of CID was varied from 2-10 µM. The datasets were fitted to a two-state function for bimolecular association to obtain estimates of the association rate constants (*kon*)^73^. (d) Observed rate constants from stopped flow displacement experiments for NCBDD/P Trp variants and NCBDD/P in complex with CID1R. The NCBDD/P variant was used to displace the different NCBDD/P Trp variants from the protein complexes, while NCBDD/P was used to displace the NCBDD/P /CID1R complex. (e) An example of an ITC binding isotherm where CID1R was titrated into NCBDD/P. All ITC measurements were conducted in the same experimental buffer as above and at 25 °C. Fitting to a one-site two-state binding model yielded the following parameters: *KD* = 2.9 ± 0.1 µM and n= 0.8 ± 0.007. (f) The affinities measured by ITC for CID1R binding to NCBDD/P, NCBDD/P and NCBDD/P. The error bars denote the standard error from the fit to a two-state function. (g) The fitted parameters for the NCBDD/P Trp variants and NCBDD/P^ML^ from the different experiments.

**Figure S2.**
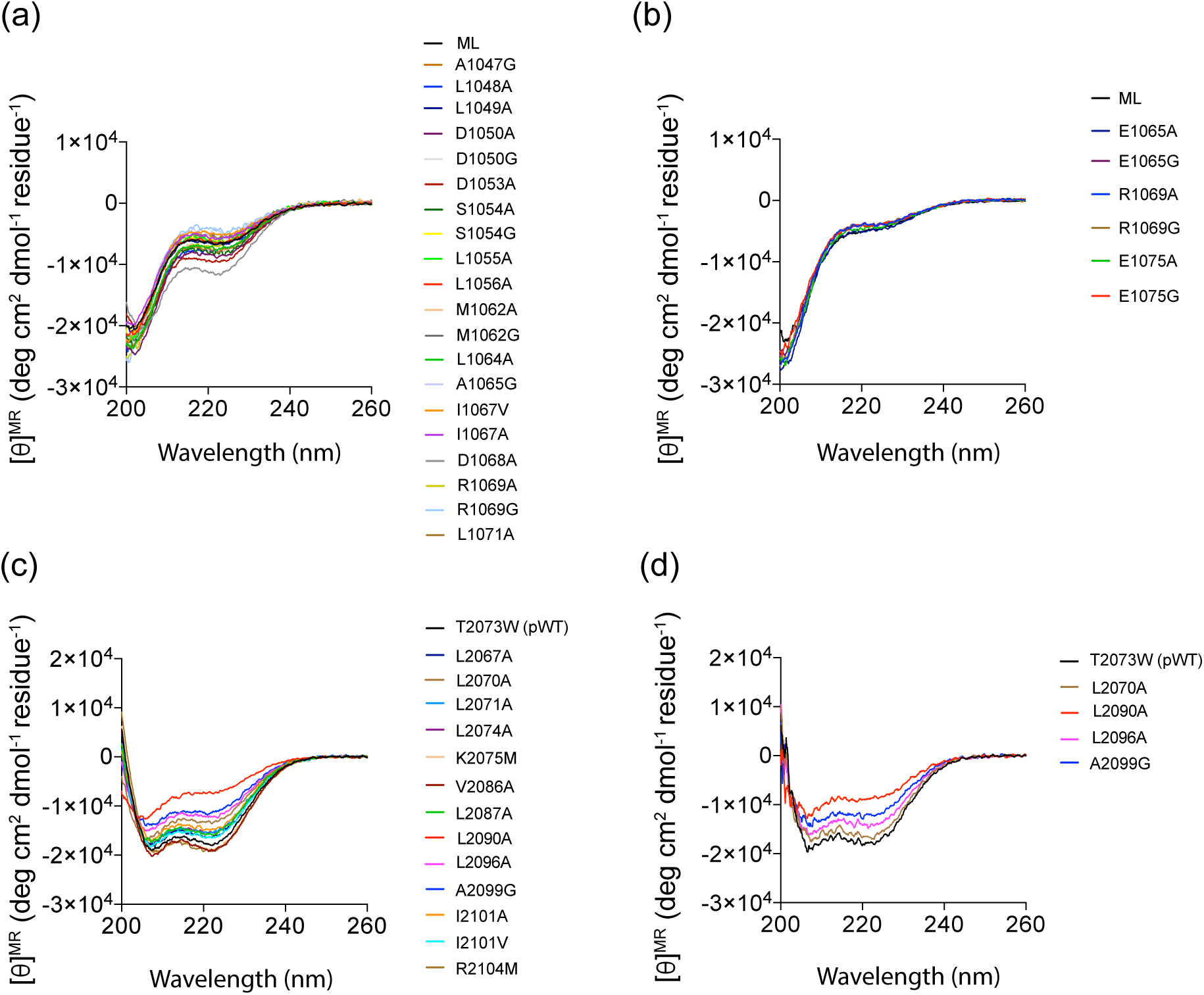
CD spectra of all NCBD and CID variants. All CD spectra were recorded in 20 mM sodium phosphate pH 7.4, 150 mM NaCl at 4°C unless stated otherwise. (a) The CD spectra of all CID1R variants. (b) The CD spectra for all CIDHuman variants. (c) The CD spectra of all NCBDD/P variants. (d) The NCBDD/P variants L2070A, L2090A, L2096A and A2099G displayed substantially lower helical content as compared to NCBDD/P and the CD spectra of these variants were therefore recorded in buffer supplemented with 0.7 M TMAO. Supplementation with TMAO increased helical content of some variants.

**Figure S3.**
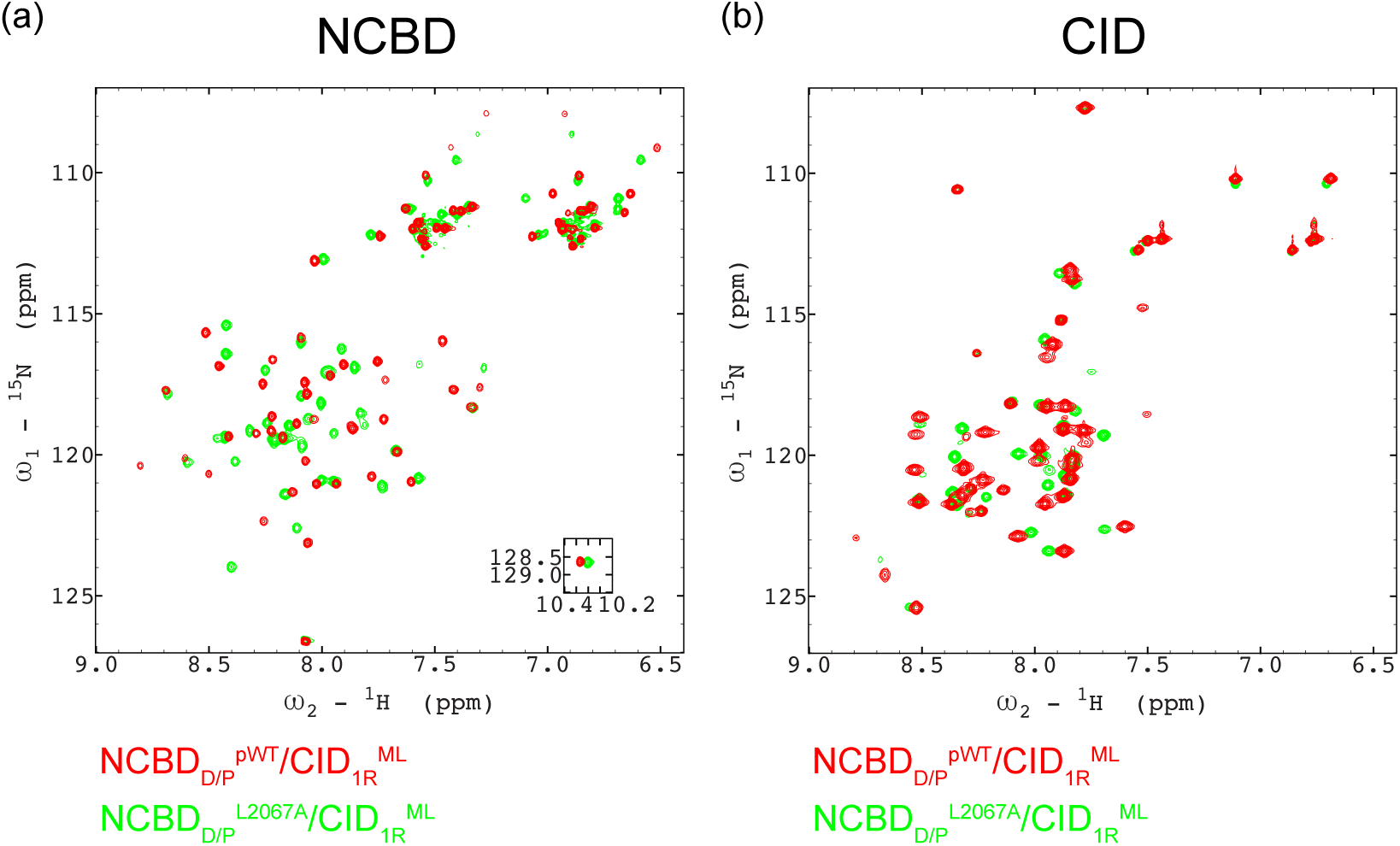
^1^H-^15^N-HSQC spectra for bound CID and NCBD domains. (a) Unlabeled CID1R^ML^ bound to ^15^N-labeled NCBDD/P^pWT^ (red) and ^15^N-labeled NCBDD/P^L2067A^ (green). The inset shows the tryptophan side chain peak. (b) ^15^N-labelled 1R CID1R^ML^ bound to unlabeled NCBDD/P^pWT^ (red) and NCBDD/P^L2067A^ (green).

**Figure S4.**
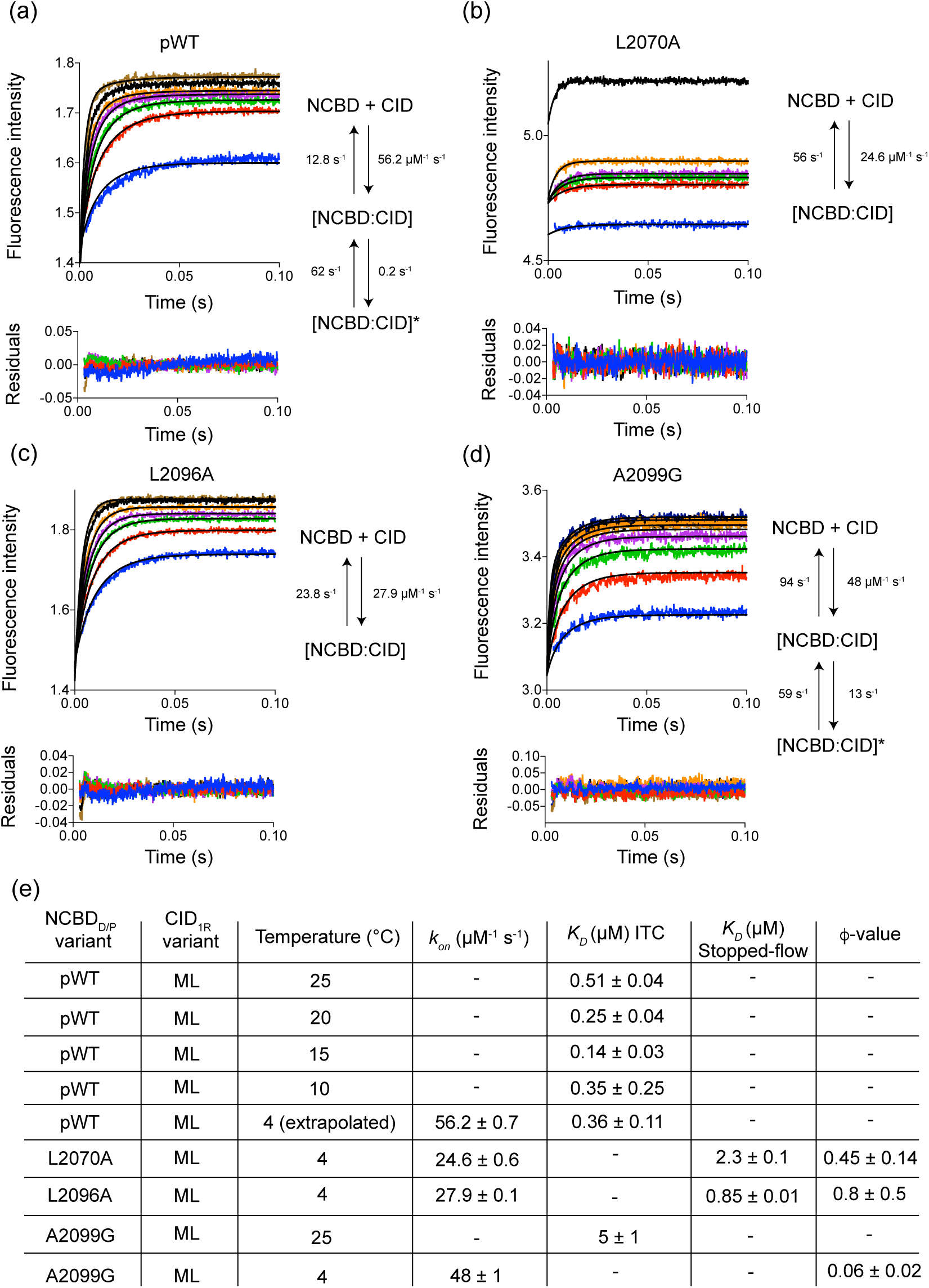
Stopped flow kinetic experiments in presence of 0.7 M TMAO. The stopped-flow binding kinetic experiments were conducted in 20 mM sodium phosphate pH 7.4, 150 mM NaCl, 0.7 M TMAO and the measurements were recorded at 4 °C. (a) The binding kinetics of NCBDD/P in complex with CID1R were biphasic in presence of 0.7 M TMAO and the figure shows the fit to an induced fit model along with the fitted microscopic rate constants. (b) The binding kinetics of NCBDD/P^L2070A^ and CID1R was monophasic and was globally fitted to a two-state binding mechanism. (c) The binding kinetics of NCBDD/P and CID1R was monophasic and the figure shows the fit to a two-state binding model. (d) The binding kinetics of NCBDD/P in complex with CID1R was biphasic in the presence of 0.7 M TMAO and was fitted to an induced fit model. (e) A table with kinetic and thermodynamic parameters obtained in stopped-flow and isothermal titration calorimetry (ITC) experiments for the protein complexes. The errors are Standard Errors from fitting a two-state model for binding. The error of the extrapolated affinity at 4°C for the NCBDD/P was estimated to be 30 %.

**Figure S5.**
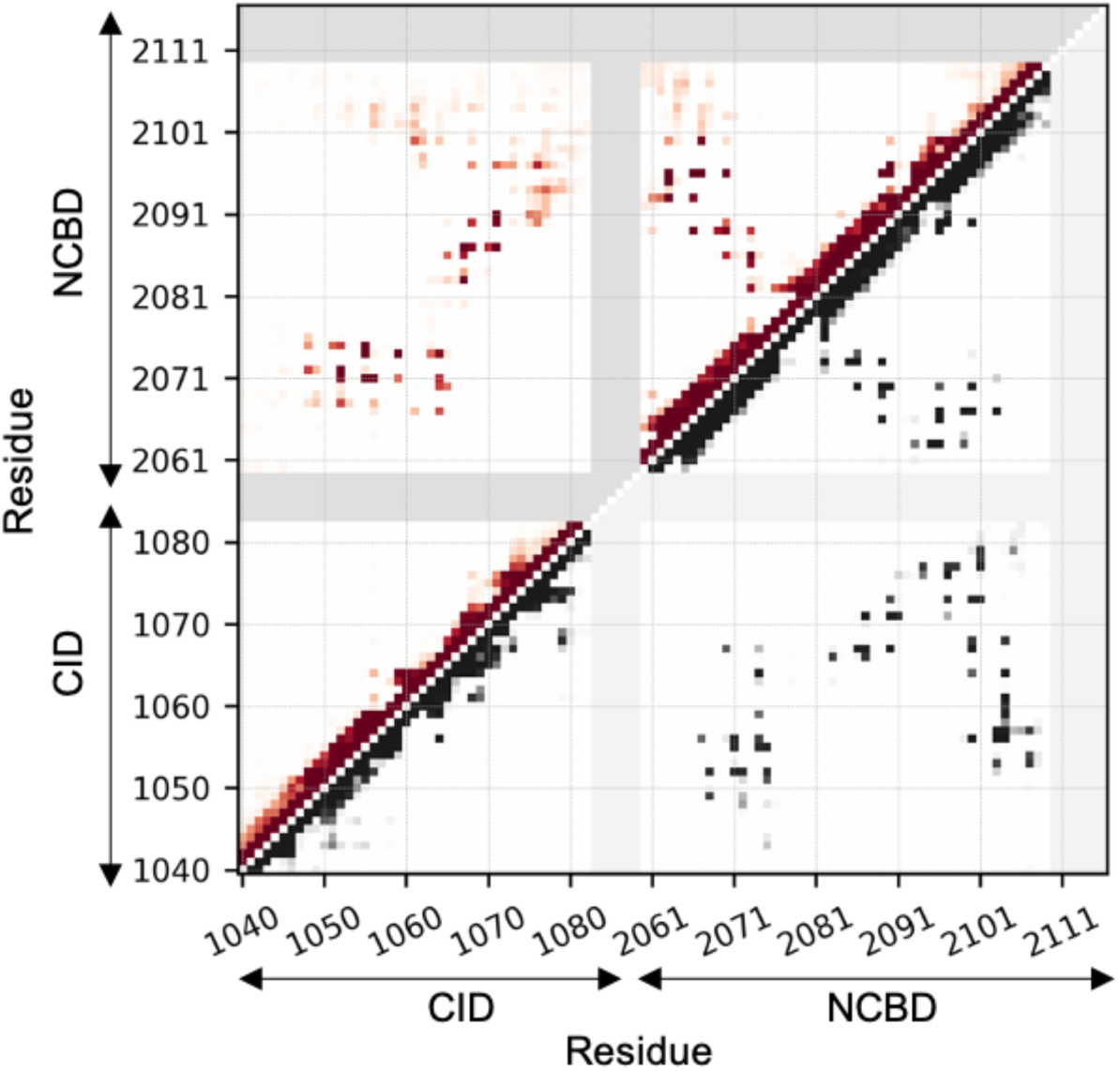
Map representing the contact probability between each pairs of residues in the ancestral native state (lower right, gray) and in the ancestral TS (upper left, red) ensembles. Probability goes from 0 (white) to 1 (dark gray/red); regions, involving residues which are present in the human but not the ancestral complex, are shaded with gray to be consistent with Figure 3 in the main text.

**Figure S6.**
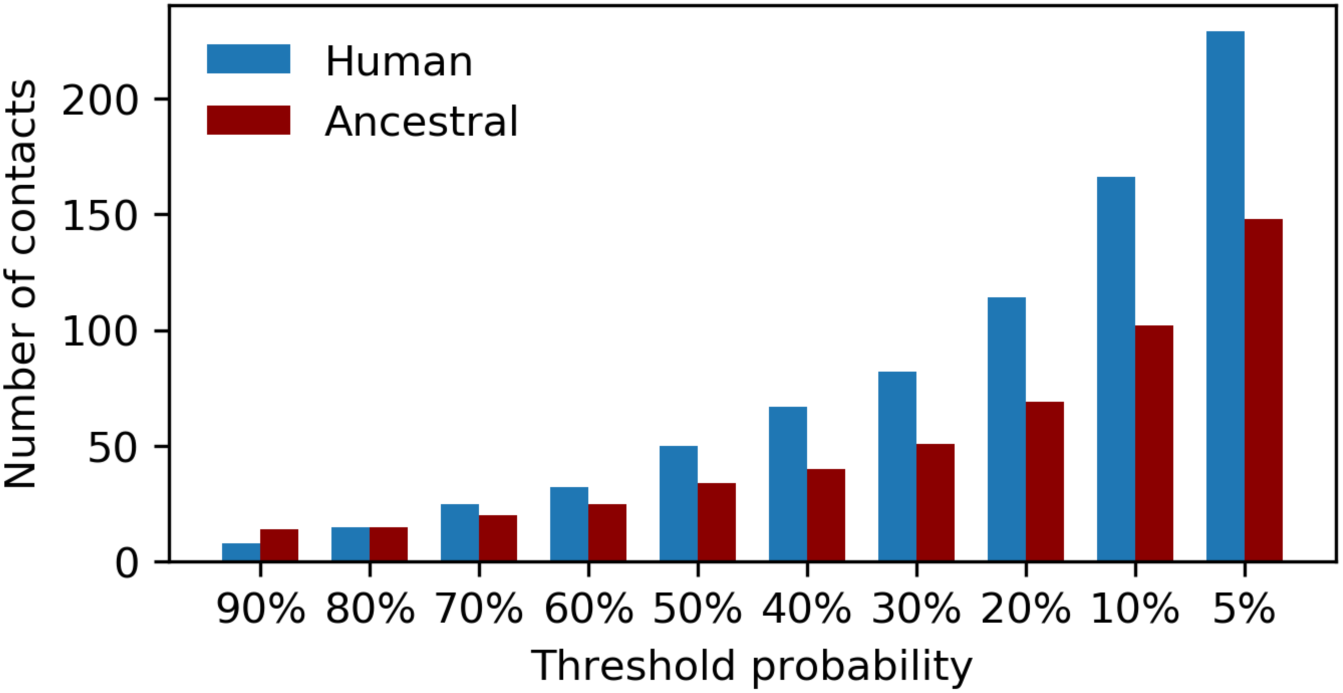
Number of intermolecular residue-residue contacts present in the human/ancestral TS ensemble with a probability higher than a threshold probability.

**Table S1.**
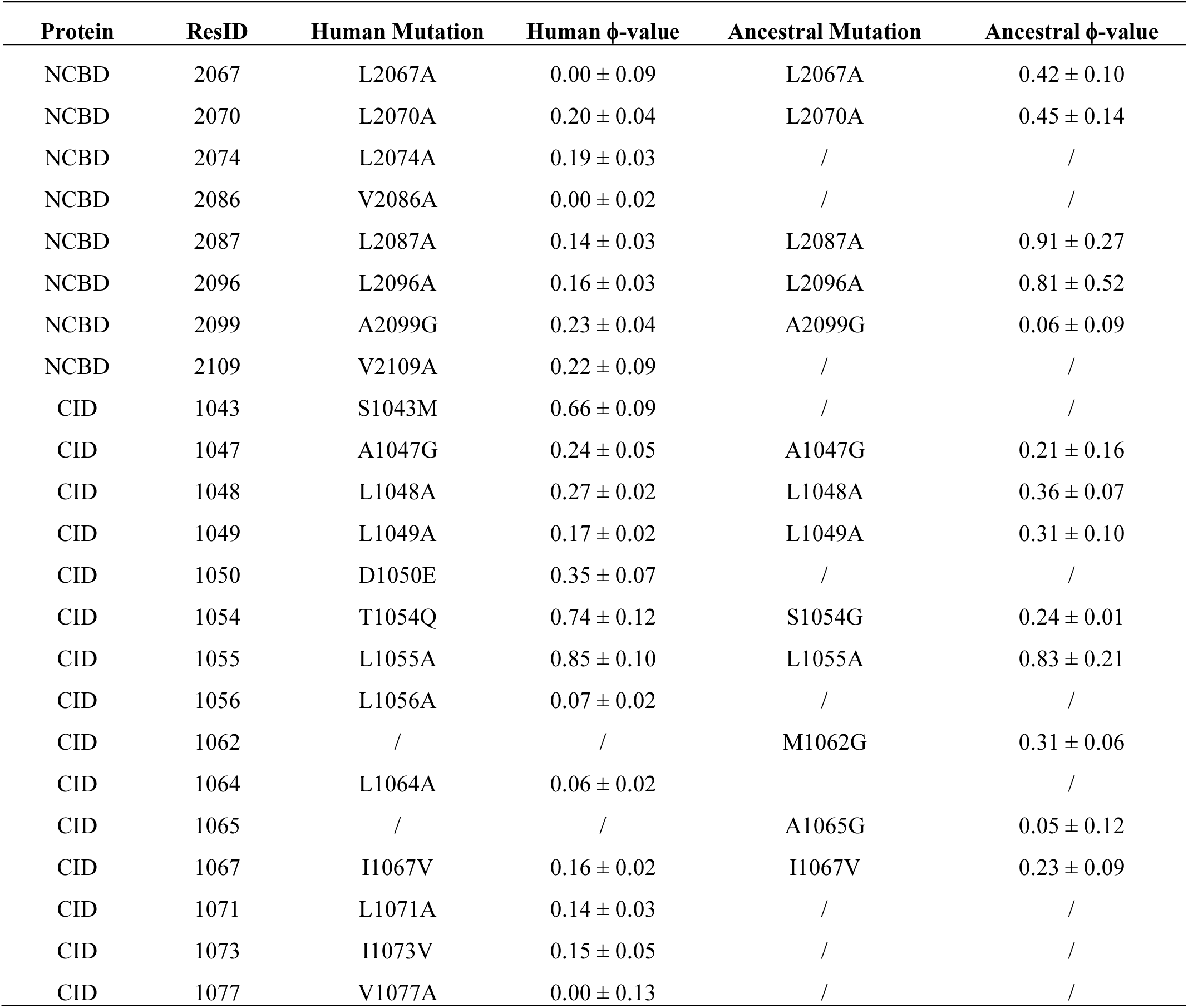
**ϕ-values used in ancestral and human TS simulations.** List of *ϕ*-values used as restrains in MD simulations for the determination of the ancestral and human TS ensembles.

